# A probabilistic functional atlas of human occipito-temporal visual cortex

**DOI:** 10.1101/2020.01.22.916239

**Authors:** Mona Rosenke, Rick van Hoof, Job van den Hurk, Kalanit Grill-Spector, Rainer Goebel

## Abstract

Human visual cortex contains many retinotopic and category-specific regions. These brain regions have been the focus of a large body of functional MRI research, significantly expanding our understanding of visual processing. As studying these regions requires accurate localization of their cortical location, researchers perform functional localizer scans to identify these regions in each individual. However, it not always possible to conduct these localizer scans. Here, we developed and validated a functional region of interest atlas of early visual and category-selective regions in human ventral and lateral occipito-temporal cortex. Results show that for the majority of fROIs, cortex-based alignment results in lower between-subject variability compared to nonlinear volumetric alignment. Furthermore, we demonstrate that (1) the atlas accurately predicts the location of an independent dataset of ventral temporal cortex ROIs and other atlases of place-selectivity, motion-selectivity, and retinotopy. Next, (2) we show that the majority of voxel within our atlas are responding mostly to the labelled category in a left-out subject cross-validation, demonstrating the utility of this atlas. The functional atlas is publicly available (download.brainvoyager.com/data/visfAtlas.zip) and can help identify the location of these regions in healthy subjects as well as populations (e.g. blind people, infants) in which functional localizers cannot be run.

## Introduction

Human visual cortex extends from the occipital lobe to the posterior parietal and temporal lobes, containing more than two dozen visual areas. Early and intermediate visual areas are typically defined by their representation of the visual field, where each visual area contains a topographic (retinotopic; Engel et al., 1994; Sereno et al., 1995) representation of the entire visual field across both hemispheres (referred to as a visual field map, Arcaro et al., 2009; Wandell et al., 2005; Wandell and Winawer, 2011; Wang et al., 2014). Higher visual areas are typically defined by their function and stimulus selectivity rather than the representation of the visual field. This includes preference to visual attributes such as motion (Sereno et al. 1995), shape (Malach et al. 1995; Grill-Spector et al. 1998), or color (Lafer-Sousa et al. 2016), as well as preference for certain visual stimuli over others. A well-documented characteristic of higher-level regions in ventral and lateral occipito-temporal cortex are regions that respond preferentially to ecologically-relevant stimuli such as faces (Kanwisher et al. 1997), places (Aguirre et al. 1998; Epstein and Kanwisher 1998), bodies (Downing et al. 2001; Peelen and Downing 2005), and words (Cohen et al. 2000) compared to other stimuli. These regions are referred to as category-selective regions.

To elucidate neural mechanisms of visual processing and perception, a central goal in neuroscience is to understand the function and computation in each of these regions. Indeed, tens of thousands of papers have investigated visual processing in specific visual areas, from visual field maps to category-selective regions. For example, according to google scholar, more than 7575 studies cite the study that discovered the fusiform face area (Kanwisher et al. 1997). The first step in this scientific endeavor is the identification of each visual region in each brain. The standard approach is to perform an independent scan, such as retinotopic mapping (Engel et al. 1997) or a functional localizer scan, in each individual to identify the relevant region of interest (ROI, Kanwisher et al. 1997; Saxe et al. 2006). Then, the main experiment of interest is performed, and the data are analyzed within the ROI identified using the independent scans. The ROI approach is advantageous for four reasons: (1) it allows hypothesis driven comparisons of signals within independently-defined regions of interest across many different conditions, (2) it increases statistical sensitivity in multi-subject analyses (Nieto-Castañón and Fedorenko 2012), (3) it reduces the number of multiple comparisons present in whole-brain analyses (Saxe et al. 2006), and (4) it identifies ROIs in each participant’s native brain space.

Nevertheless, there are also several limitations to the independent localizer approach. First, it is not always possible to obtain an independent localizer scan. This is especially the case in patient populations, for example in the congenitally blind (Mahon et al. 2009; Bedny et al. 2011; Striem-Amit, Cohen, et al. 2012; van den Hurk et al. 2017) or individuals with visual agnosia/prosopagnosia (Schiltz and Rossion 2006; Steeves et al. 2006; Sorger et al. 2007; Barton 2008; Gilaie-Dotan et al. 2009; Susilo et al. 2015). Second, performing a localizer scan before each experiment is costly in terms of scanning time, as well as mental effort and attention resources of the participant. The latter can result in fatigue during the main experiment of interest, leading to lower quality data. Third, as localizer scans are typically conducted in a subject-specific manner, and researchers vary in the manner they define the ROIs (e.g. whether smoothing was employed, if they use anatomical constraints, what thresholding methods were employed), it is hard to assess variability between participants and across studies.

To overcome these limitations, progress in the field of cognitive neuroscience has led to the development of cortical atlases, which allow localization of visual areas in new subjects by leveraging ROI data from an independent set of typical participants (Frost and Goebel 2012; ventral-temporal cortex category selectivity: Julian et al. 2012; Engell and McCarthy 2013; Zhen et al. 2017a; Weiner et al. 2018; visual field maps: Benson et al. 2012; Wang et al. 2014; motion-selective hMT: Huang et al. 2019; multimodal parcellation: Glasser et al. 2016; cytoarchitectonic parcellation of ventral visual cortex: Rosenke et al. 2018). In addition to providing independent means to identify ROIs, this approach enables quantification of between-subject variability. Further, the process of atlas creation also enables measuring the prevalence and robustness of each ROI across participants. Presently, atlases for the human visual system include atlases of visual field maps (Benson et al. 2012, 2014; Wang et al. 2014; Benson and Winawer 2018), and atlases of cytoarchitectonically-defined areas (Amunts et al. 2000; Rottschy et al. 2007; Caspers et al. 2013; Kujovic et al. 2013; Lorenz et al. 2015; Rosenke et al. 2018). However, presently, there is no atlas of the full extent of visual category-selective regions in occipito-temporal cortex, or atlases that include both visual regions that are defined retinotopically as well as from stimulus selectivity. To fill this gap in knowledge, in the present study we: (a) develop a functional atlas of category-selective visual cortex, (b) quantify inter-subject variability of category-selective regions in visual cortex, and (c) validate our approach by using the same procedure to define retinotopic regions and hMT+, which also allows us to compare our definitions to existing atlases. To generate the visual functional atlas (visfAtlas), 19 participants (10 female) underwent the following functional scans: (i) a localizer experiment to identify word, body, face, body, and place-selective regions in lateral occiptio-temporal (LOTC) and ventral temporal cortex (VTC), (ii) a visual field mapping experiment to delineate early visual cortex (V1-V3), and (ii) a motion localizer to identify hMT+. We identified each ROI in each participant’s brain. We then used a leave-one-out cross-validation (LOOCV) approach and two anatomical alignment methods: (i) nonlinear volume-based alignment (NVA) and (ii) cortex-based alignment (CBA), to evaluate the accuracy of the atlas in predicting ROIs in new participants. The resulting visfAtlas is available with this paper in BrainVoyager (www.brainvoyager.com) and FreeSurfer (www.surfer.nmr.mgh.harvard.edu) file formats for cortical surface analyses, as well as in nifti format for volumetric analysis (download.brainvoyager.com/data/visfAtlas.zip).

## MATERIALS AND METHODS

### Participants

To obtain functional data, a total number of 20 participants (average age 30 ± 6.61) were recruited at Maastricht University but one subject’s functional MRI (fMRI) scans were excluded from further analysis due to self-reported lack of attention on the stimuli and intermittent sleep. Two participants were left-handed, and the sample consisted of 10 women and 9 men. All participants were healthy with no history of neurological disease and had normal or corrected-to-normal vision. Written consent was obtained from each subject prior to scanning. All procedures were conducted with approval from the local Ethical Committee of the Faculty of Psychology and Neuroscience.

### Data acquisition

Participants underwent one scanning session of 1 hour at a 3T Siemens Prisma Fit (Erlangen, Germany). First, a whole brain, high resolution T1-weighted scan (MPRAGE) was acquired (repetition time/echo time = 2250/2.21 ms, flip angle = 9 °, field of view = 256 x 256 mm, number of slices = 192, 1 mm isovoxel resolution). Following that, six functional runs were acquired using a T2*-weighted sequence with the following parameters: repetition time/echo time = 2000/30 ms, flip angle = 77 °, field of view = 200 x 200 mm, number of slices = 35, slice thickness = 2 mm, in-plane resolution = 2 × 2 mm. fMRI included (i) three scans of the functional localizer (fLoc; Stigliani et al. 2015) (ii) two scan of an hMT+ localizer, and (iii) one scan of retinotopic mapping. Maximal diameter of the visual stimuli ranged from 30°-36° in the fMRI experiments. Details for each localizer can be found in the section below.

### Visual localizers

#### Category-selective regions in ventral temporal cortex and lateral occipito-temporal cortex

In order to identify category-selective regions that respond preferentially to characters (pseudowords, numbers), bodies (whole bodies, limbs), places (houses, corridors), faces (child, adult) and objects (cars, instruments), we used stimuli included in the fLoc functional localizer package (Stigliani et al. 2015). Eight stimuli of one of the five categories were presented in each miniblock design, each miniblock holding a duration of 4 seconds. To assure participant’s attention, they were asked to perform an Oddball task, indicating with a button press when they saw a scrambled image instead of one of the categories. Each run consisted of 150 volumes, and each subject underwent three runs.

#### hMT+

To localize the motion-selective area in middle temporal cortex (hMT+, Dumoulin et al., 2000; Zeki et al., 1991), we used stimuli as in Emmerling et al. (2016) and Zimmermann et al. (2011), which were based on Huk et al. (2002). During the first 5 volumes participants were presented with a fixation dot in the center of the screen. In the following blocks, moving and stationary dot patterns alternated while the participants fixated on the fixation dot at the center of the screen. Moving dot blocks were 18 seconds long, while stationary blocks had a duration of 10 seconds. The active screen filled with dots was circular. In total, each run consisted of 12 blocks of moving dots and 12 blocks of stationary dots. Black dots on a gray background traveled towards and away from the fixation point (speed = 1 pixel per frame, dot size = 12 pixels, number of dots = 70). In different blocks, dots were presented either in the center of the screen, in the left visual hemifield, or in the right visual hemifield. Stationary blocks were in the same three locations. The order of blocks was fixed (center moving, center static, left moving, left static, right moving, right static). Each subject underwent two hMT+ localizer runs.

#### Early visual cortex

We ran one visual retinotopic mapping run that consisted of 304 volumes (TR = 2s). In the first 8 volumes a fixation dot was presented, followed by a high-contrast moving bar stimulus (1.33° wide) revealing a flickering checkerboard pattern (10 Hz). The checkerboard pattern varied in orientation and position for 288 volumes, concluding the run with 8 volumes of fixation dot presentation. The fixation was presented during the entire run and changed color at random time intervals. To keep participants’ motivation and attention they were asked to count these color changes. The bar stimulus moved across the visual field in 12 discrete steps and remained at each position for 1 TR. The 12 different stimulus positions were randomized within each bar orientation. Each combination of orientation (4) and direction (2) represented one cycle. These eight different cycles were repeated three times in random order throughout the run (Senden et al., 2014).

### Preprocessing

If not stated otherwise, data were preprocessed and analyzed using BrainVoyager 20.6 (Brain Innovation, Maastricht, The Netherlands). Anatomical data were inhomogeneity corrected and transformed to Talairach space (TAL, Talairach and Tournoux, 1988) by identifying the anterior commissure (AC) and posterior commissure (PC) and fitting the data to TAL space. Functional data were slice scan time corrected, motion corrected with intra-run alignment to the first functional run to account for movement between runs, and high-pass filtered (3 cycles). Next, the preprocessed functional data were co-registered to the inhomogeneity corrected anatomical image. Using the anatomical transformation files, all functional runs were normalized to TAL space. Based on the normalized anatomical data, we segmented the grey-white matter boundary for each brain and created a cortical surface. Next, the volumetric functional data were sampled on the cortical surface incorporating data from −1 to +3 mm along the vertex normals. Ultimately, we computed two general linear models (GLM), one for the three localizer runs for category-selective regions in ventral temporal cortex, and one for the hMT+ localization.

### Regions of interest

All ROIs where manually defined in individual subjects on their cortical surface reconstruction in BrainVoyager. For volumetric alignment and atlas generation, surface regions were transformed to volumetric regions by expanding them (−1 to +2 mm) along the vertex normals of the white-gray matter boundary. The final atlas includes all regions that could be defined in more than 50% of the subjects (N*≥*10, see Table 1 for number of subjects per atlas ROI).

**Table 1.** Nomenclature for functional regions-of-interest (fROIs) and number of subjects per fROI. Nomenclature: Each category-selective functional activation cluster can be described by functional category or anatomical location. In this article we describe category-selective ROIs using the anatomical nomenclature and provide this table as a reference. Functional abbreviations are as followed: *FFA*: fusiform-face area, *FBA*: fusiform-body area, *EBA*: extrastriate body area, *VWFA*: visual word form area, *PPA*: parahippocampal place area, *hMT*: human middle-temporal (cortex). Number of identified ROIs per hemisphere (N LH/N RH): Due to individual-subject variability and using a strict statistical threshold (*t*>3, vertex level), not every fROI was identified in all participants in both hemispheres. fROIs that were defined in more than half the participants (N≥10) were included in the atlas. Areas that were not included are indicated in gray subject counts. The last column, *N,* indicates the number of subjects in which a given fROI could be identified in at least one hemisphere. Abbreviations: *LH*: left hemisphere, *RH*: right hemisphere.

| ROI | Functional nomenclature | N (LH) | N (RH) | N |
| --- | --- | --- | --- | --- |
| mFus - faces | FFA-2 | 13 | 15 | 18 |
| pFus - faces | FFA-1 | 17 | 15 | 19 |
| IOG - faces | - | 15 | 15 | 18 |
| OTS - bodies | FBA | 14 | 13 | 17 |
| ITG - bodies | EBA | 17 | 17 | 19 |
| MTG - bodies | EBA | 16 | 15 | 18 |
| LOS - bodies | EBA | 15 | 16 | 19 |
| pOTS - characters | VWFA-1 | 16 | 5 | 17 |
| IOS - characters | - | 11 | 1 | 11 |
| TOS - places | - | 9 | 12 | 13 |
| CoS - places | PPA | 18 | 19 | 19 |
| hMT - motion | hMT | 18 | 16 | 19 |
| V1d |  | 19 | 19 | 19 |
| V2d |  | 19 | 19 | 19 |
| V3d |  | 14 | 17 | 17 |
| V1v |  | 19 | 19 | 19 |
| V2v |  | 19 | 19 | 19 |
| V3v |  | 19 | 19 | 19 |

#### Retinotopic areas in occipital cortex

Visual field maps were determined for each subject based on an isotropic Gaussian population receptive Field (pRF) model (Dumoulin and Wandell 2008; Senden et al. 2014). The obtained pRF maps estimating the location and size of a voxel pRF were used to calculate eccentricity and polar angle maps. The polar angle maps were projected onto inflated cortical surface reconstructions and used to define six topographic regions in occipital cortex (V1d, V2d, V3d and V1v, V2v, V3v, where d = dorsal and v = ventral) by identifying the reversals in polar angle representation at the lower vertical meridian (LVM), upper vertical meridian (UVM) or horizontal meridian (HM; DeYoe et al., 1996; Engel et al., 1997; Sereno et al., 1995). We did not define visual areas beyond V3d and V3v as visual field maps using the single run retinotopic mapping paradigm were noisy beyond V3.

#### Ventral and lateral category-selective areas

Each category (e.g. faces) was contrasted against the mean of all other categories to identify vertices that displayed a preference for the given category. Then we followed a two-step approach to define ROIs: First, for all categories we selected a statistical threshold of *t* = 3 for a whole brain map. Based on the thresholded activation map we identified ROIs in anatomically plausible locations (see details for each region below). Furthermore, in the case of an activation cluster transitioning into an adjacent one of the same visual category, we divided those clusters into separate ROIs by following the spatial gradient of *t*-values and separating the two areas at the lowest t-value. Based on insufficient activation pattern found for the ‘objects’ category, we dismissed that category from further analysis.

Face-selective regions (faces > all others) were identified in the mid lateral fusiform gyrus (mFus) and posterior lateral fusiform gyrus (pFus), which correspond to the fusiform face area (Kanwisher et al. 1997), as well as on the inferior occipital gyrus (IOG). Body-selective regions (bodies > all others) were observed in ventral temporal cortex on the occipital temporal sulcus (OTS), also known as fusiform body area (FBA, Peelen et al., 2009; Schwarzlose, 2005) and in lateral occipital cortex. There, we identified three different regions (Weiner and Grill-Spector 2011) together forming the extrastriate body area (Downing et al. 2001), one anterior of hMT+ on the middle temporal gyrus (MTG), one posterior of hMT+ on the lateral occipital sulcus (LOS), and one ventral to hMT+, on the inferior temporal gyrus (ITG). Place-selective regions (places > all others) were observed in ventral temporal cortex on the collateral sulcus (CoS), corresponding to the parahippocampal place area (PPA, Epstein and Kanwisher, 1998), and on the transverse occipital sulcus (TOS, Hasson et al., 2003). Character-selective regions (characters > all others) were identified in the posterior occipital temporal sulcus (pOTS) and a left-lateralized region in the mid occipital temporal sulcus (mOTS). Furthermore, we identified one character-selective regions in the inferior occipital sulcus (IOS). In the following, we will refer to each ROI by its anatomical nomenclature, as described in Stigliani et al. (2015). For reference, Table 1 provides an overview about each ROI’s anatomical as well as functional name.

#### hMT+

Motion selective regions were identified by contrasting left, right and central visual field motion conditions vs. the equivalent stationary conditions and using a thresholded statistical map with a minimum *t*-value of 3. Two subjects only showed functional activation for the contrasts at a *t*-value of 2.5 in one hemisphere, which we allowed for these subjects. hMT+ was consistently located in the posterior inferior-temporal sulcus (pITS).

### Visual functional atlas (visfAtlas) generation

After ROIs were defined for each subject in each subject’s space, we utilized two normalization techniques to bring the data into a common space: (1) nonlinear volumetric alignment (NVA) for volume and (2) cortex-based alignment (CBA) for surface space. Furthermore, as it is common that not every ROI can be identified in each of the subjects, we decided that an ROI had to be present in more than 50% of the subjects (N > 10) to be considered for a group atlas. The ROIs which were ultimately used for the group atlases and in how many subjects they were defined can be found in Table 1.

#### Nonlinear-volumetric alignment (NVA)

First, surface regions that were defined on each subject’s cortical surface were mapped to volumetric regions by expanding them (−1 to +2 mm) along each vertex normal of the white-gray matter boundary. Second, the volumetric regions were transformed back to native ACPC space.

Next, the individual brains were registered to the MNI152 group average brain using the Advanced Normalization Tools (ANTS; https://sourceforge.net/projects/advants/). Finally, the resulting nonlinear transformation matrices were used to warp the functionally-defined regions of interest (fROIs) into the same orientation and reference frame. The specific code for the affine volume registration and nonlinear transformation can be found here: download.brainvoyager.com/data/visfAtlas.zip. The resulting NVA-aligned regions were further processed in NifTi format using MATLAB 2014b and 2019a (www.mathworks.com), see details below.

#### Cortex-based alignment (CBA)

To generate a surface group average brain of the subjects, we used cortex-based alignment (CBA) to generate a dynamic average (subsequently called BVaverage, publicly available at download.brainvoyager.com/data/visfAtlas.zip and usable as surface template for future studies). CBA was performed for both hemispheres separately after inflation to a sphere with overlaid curvature information at various levels of resolution (Goebel et al. 2006; Frost and Goebel 2012). First, during a rigid alignment, the spheres of each subject’s hemisphere was rotated along three dimensions to best match the curvature pattern of a randomly chosen target hemisphere. The lower the variability between the two folding curvature patterns, the better the fit after rigid sphere rotation. Following the rigid alignment for all subjects, a non-rigid CBA was performed. Curvature patterns of each subject were used in four different levels of anatomical detail. Starting from low anatomical detail, each subject’s hemisphere was aligned to a group average out of all subjects. During this process, the group average was dynamically updated to most accurately average all hemispheres. This sequence was repeated for all levels of curvature detail, until the group average was updated based on the highest level of anatomical detail per subject. During the alignment, we (1) derived a group average for each hemisphere (BVaverage), as well as (2) a transformation indicating for each vertex on a single-subject cortical surface where it maps to on the group average. These transformation files were then used to map each individual subject’s fROIs to the BVaverage.

#### Probabilistic maps for occipitotemporal cortex in volume and surface space

We generated probabilistic maps of all regions after NVA as well as CBA, where each of the following was done in both group spaces: after individual subject fROIs were projected to the MNI152 and BVaverage, respectively, each group fROI was defined. For each voxel/vertex of a group fROI, the number of subjects sharing that voxel/vertex in the fROI was divided by the total number of subjects of the fROI 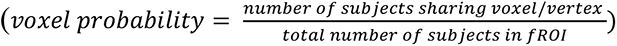. Thus, a value of 0 at a vertex in the group fROI indicates a vertex did not belong to that fROI in any subject, a value of .5 means that it belonged to the fROI in half the subjects, a value of 1 indicates that it belonged to that functional region in the entire study population (Fig. 1).

**Figure 1.**
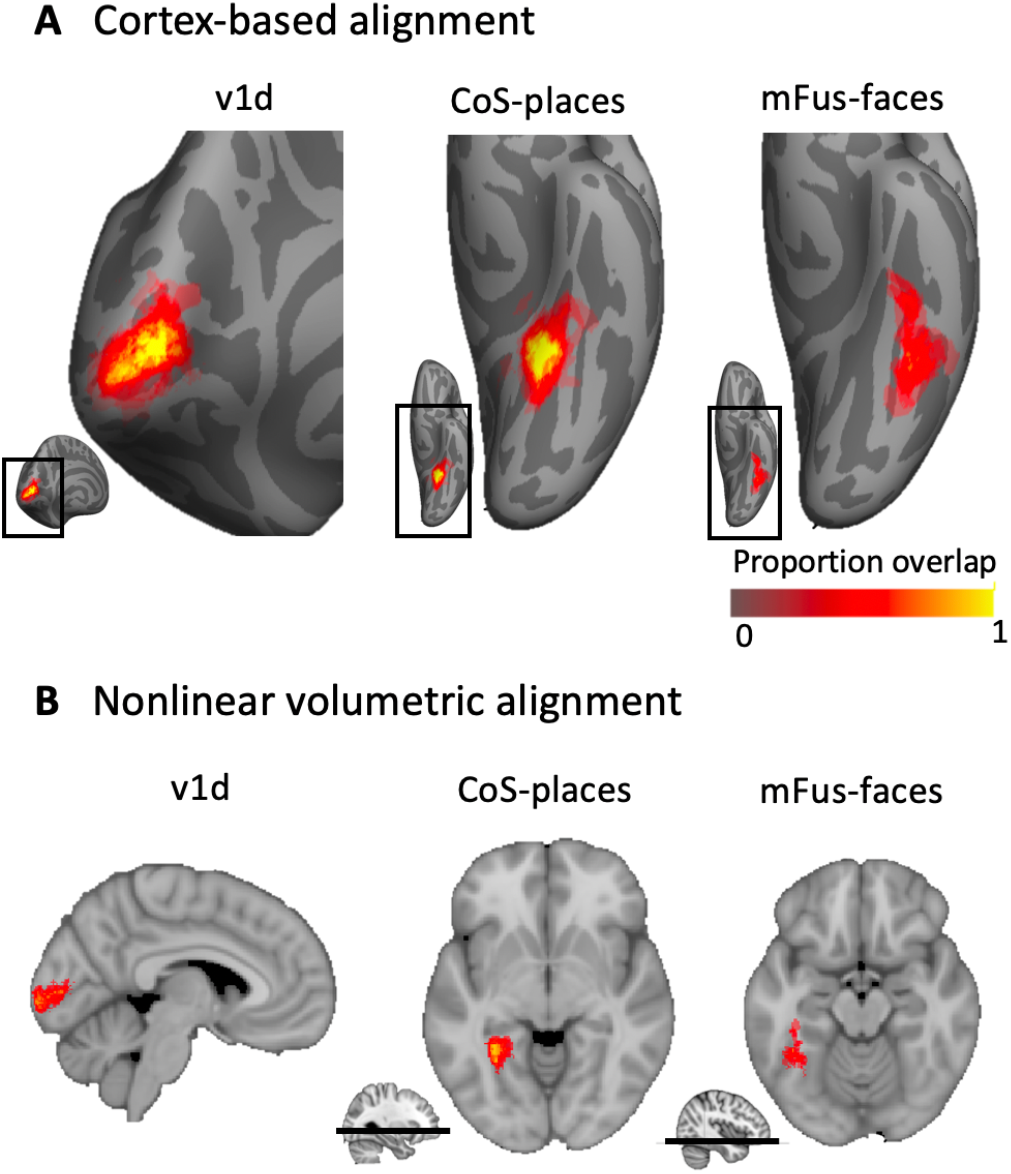
Example probabilistic group maps in the left hemisphere after two brain alignments. **(A)** Three example regions-of-interest (ROIs) are displayed where the most left column, v1d, shows an early visual cortex map and the middle and right columns display two higher-order visual category-selective regions in ventral temporal cortex, Cos-places and mFus-faces. Probability values range from 0 to 1 where 0 indicates no subject at a given vertex and 1 that all subjects in the probabilistic maps shared the given vertex. mFus-faces reveals less consistency as shown by a lower percentage of yellow-colored vertices. Bottom inset displays zoomed in location of the main figure*. **(B)*** Same ROIs as in A but after nonlinear volumetric alignment (NVA). Bottom inset for CoS-places and mFus-faces indicates the location of the axial slice in the volume.

#### Cross-validated predictability estimation and atlas generation

One interesting feature of those fROIs is the possibility to serve as a prior to estimate the localization of corresponding ROIs in a new subject’s brain, eliminating the need for a dedicated localizer run in the new subject. To allow for a probabilistic estimate to find this region in a new subject, we performed an exhaustive leave-1-subject-out cross-validation analysis after the volumetric (NVA) as well as surface (CBA) alignment to establish how well our atlas can predict fROIs in new subjects. For each fold of the LOOCV, we generated a group probabilistic fROI (G) and a left-out subject’s individual fROI (I). We estimated the predictability of the group probabilistic fROI by calculating the Dice coefficient (DSC), a measure of similarity of two samples:

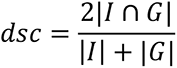

A Dice coefficient of zero indicates no predictability and a Dice coefficient of 1 indicates perfect predictability. As we did in previous work (Rosenke et al. 2018), we applied different threshold levels to the group probabilistic fROI (G) to predict the location of the left-out-subject (Fig. 2). That means we created a liberal group probabilistic fROI including each vertex that was present in at least 1 subject. Then we sequentially increased the threshold up to the most conservative threshold where all subjects had to share a voxel/vertex for it to be included in the group map. For statistical assessment, we compared Dice coefficients across the two alignment methods using a repeated measures analysis of variance (ANOVA) with individual regions as different entries, alignment method (CBA vs. NVA) as within-subject factor, and hemisphere as between-subject factor. We ran this comparison on two different thresholds: once on unthresholded group maps, and once on a threshold that produced - across regions and methods - the highest predictability. To determine this threshold, we averaged Dice coefficient values across alignment methods, hemispheres, and ROIs, resulting in one Dice coefficient per threshold level (as previously done in Rosenke et al. 2018). Comparison across thresholds revealed that a threshold of 0.2 produced the highest predictability. Additionally, we ran paired permutation tests within each region on Dice coefficient results at threshold 0.2 to establish whether the specific region showed a significant Dice coefficient for either alignment (NVA or CBA). Finally, we calculated the mean ROI surface area (in mm^2^) for each hemisphere and ROI (Fig. 3) and used a paired *t*-statistic to assess whether there was a systematic hemispheric difference in size across ROIs.

**Figure 2.**
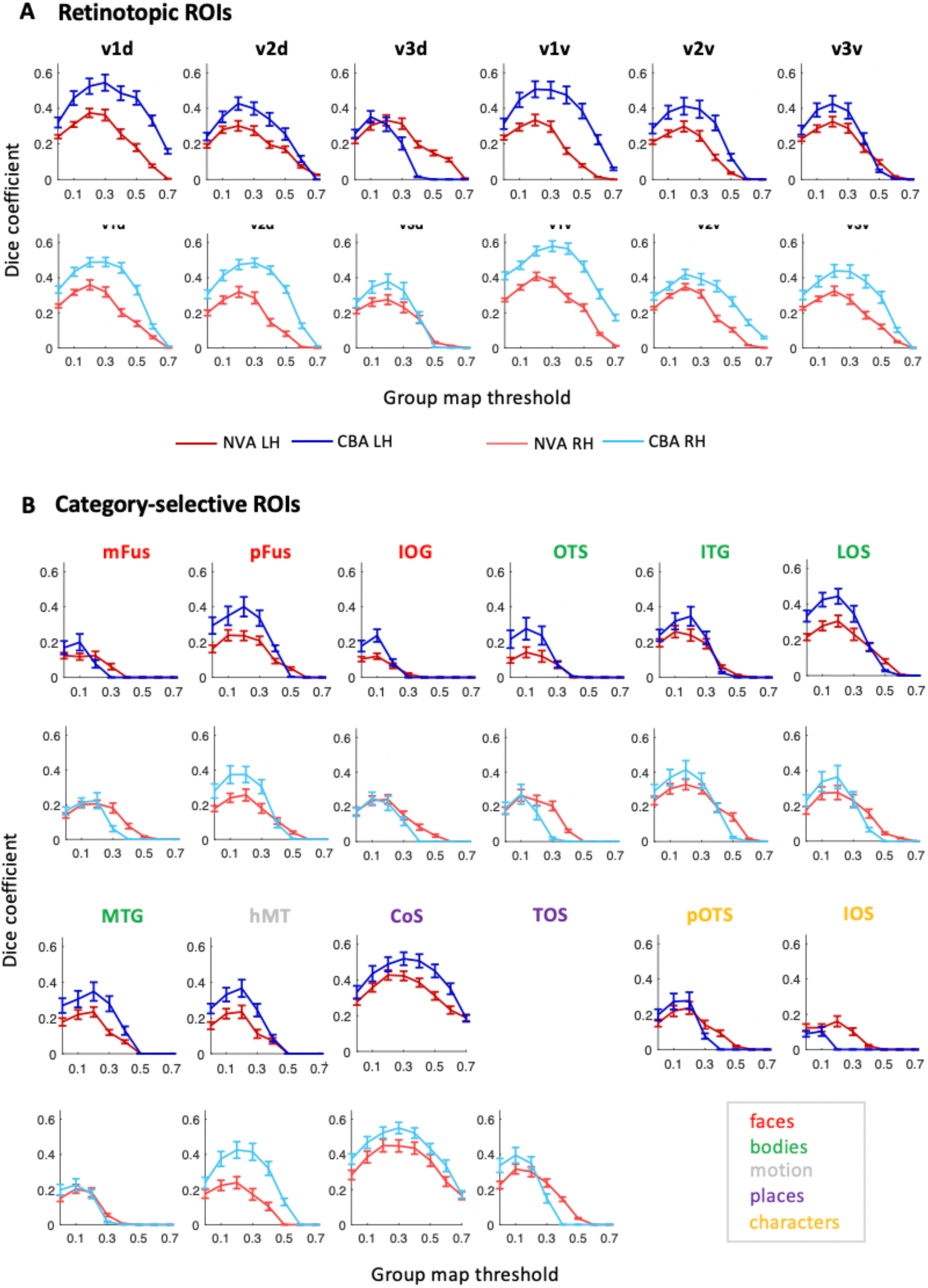
Leave-one-out cross-validation predictability analysis using the Dice coefficient (DSC) for retinotopic regions (A) and category-selective regions (B). *x-axis:* threshold of the probability map generated using N-1 subjects, y-axis: DSC. A DSC value of 1 indicates perfect overlap between the N-1 group map and the left-out subject, 0 indicates no overlap. *Blue lines*: DSC after CBA, *red lines*: DSC after NVA. Dark colors/top rows correspond to left hemisphere data, light colors/bottom rows to right hemisphere data. *Red*: face-selective ROIs, *green*: body-selective ROIs, *yellow:* character-selective ROIs, *gray:* motion-selective ROI, *error bars*: standard error (SE) across the N-fold cross-validation.

**Figure 3.**
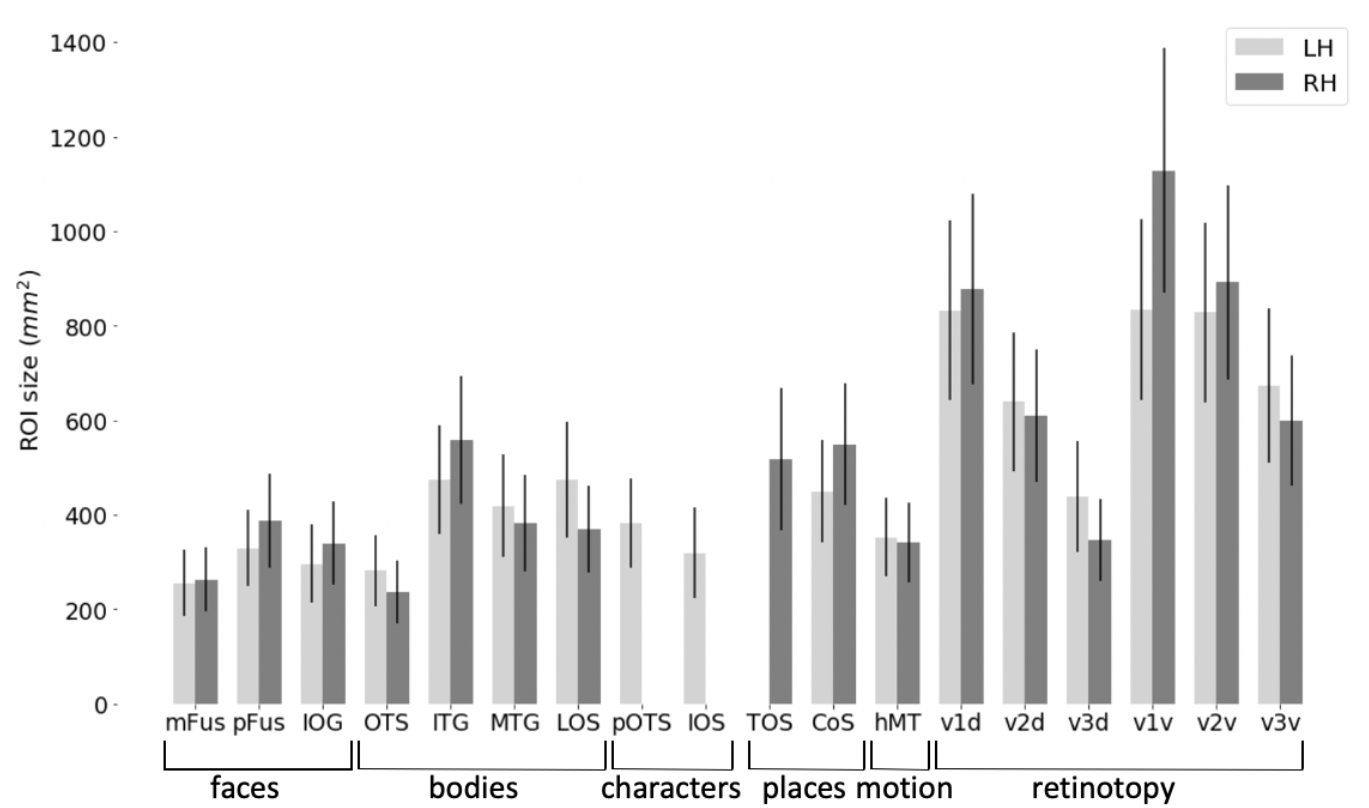
fROI size across occipito-temporal cortex. Average ROI size in surface space separately for the left hemisphere (LH, *light gray*) and right hemisphere (RH, *dark gray*). *Error bars*: standard error across subjects. Regions of X-axis are organized by category.

#### Generating a visual functional atlas (visfAtlas) by assigning each voxel and vertex to a unique <u>fROI</u>

The processes described below provide a non-overlapping tiling of the functionally defined regions in occipito-temporal cortex in surface as well as volume space (Fig. 5).

##### Cortex-based alignment

The probability maps determine the probability that each vertex belongs to a given fROI. However, it is possible that a point on the brain may belong to more than one probabilistic fROI. This overlap is more likely to occur along boundaries of neighboring functional regions. In order to assign a unique functional label to each vertex in the atlas, we generated a maximum-probability map (MPM) of each area, once in volume space (NVA) and once in surface space (CBA). Using the probabilistic fROIs, we determined which vertices were shared by more than one probabilistic fROI and assigned these vertices to a single fROI based on the area which showed the highest probability at that vertex (Eickhoff et al. 2005). In cases where two areas held the same probability value for one vertex, we averaged the probabilistic values of neighbors of that vertex for each of the fROIs. The degree of neighbors averaged was increased until the vertex had a higher probability value in one of the areas. Lastly, after all vertices were assigned in each of the MPM areas, we searched for individual vertices that were not connected to other vertices of the same ROI. We used a decision threshold where a minimum of at least one 3^rd^ degree neighbor for each vertex had to be in the same group ROI for that vertex to be part of the group ROI. In cases where single vertices where detected, they were assigned to the ROI with the second-highest probabilistic value and same-ROI vertices in the immediate neighborhood.

##### Nonlinear volume alignment

The creation of a maximum probability map in volume space was identical to that for CBA as described above, except for the neighborhood search. The neighborhood search was implemented differently as the 3D nature of the volume atlas would lead to inevitable differences in the MPM creation when compared to the surface atlas. Neighborhood search was only performed for 1 immediately adjacent voxel in all three dimensions.

### A visual functional atlas available in volume and surface space

The unique tiling of functionally defined visual regions provides a functional atlas (visfAtlas) which we make available (1) in volume space, and (2) in surface space. In addition, we make this atlas available in multiple file formats. *Volume:* we publish the volumetric visfAtlas in MNI space in BrainVoyager file format (VOI file) and NifTi format, which can be read by a variety of software packages. *Surface:* we publish the visfAtlas in file formats compatible with Brain Voyager as well as FreeSurfer. Note, however, that the surface atlases are generated slightly differently for each software. For BrainVoyager, we generated a publicly available dynamic group average brain (BVaverage, Fig. 5C) that will be available with the distributed atlas, details are described above. Since FreeSurfer (https://surfer.nmr.mgh.harvard.edu/) is commonly used with the fsaverage brain, an average surface of 39 individuals, we converted the individually defined fROIs from each subject to cortical surface space in FreeSurfer after running each subject through the recon-all pipeline. Then, we used the FreeSurfer CBA algorithm to bring each subject’s fROIs to the fsaverage space. Further processing was done as described above and the same for both software packages. All files can be downloaded here: download.brainvoyager.com/data/visfAtlas.zip.

### Evaluating whether fROI size and reproducibility are related to inter-subject consistency

There are several factors that can influence consistency across subjects. First, region of interest size has been shown to influence across-subject consistency measures using the Dice coefficient (Rosenke et al. 2018). Therefore, we determined if there is a correlation between the cross-validated Dice coefficient and average fROI surface area. Second, we established whether categories differ in reproducibility of cortical responses within a subject. We reasoned that across-subject variability cannot be expected to be lower than within-subject variability over time (reproducibility), hence it can be used as a proxy for noise ceiling. To measure reproducibility, we first defined two regions of interest, ventral temporal cortex (VTC) and lateral occipito-temporal cortex (LOTC). VTC was manually defined by tracing well known anatomical: the occipitotemporal sulcus (OTS), posterior transverse collateral sulcus (ptCoS), parahippocampal gyrus (PHG) and the anterior tip of the mid-fusiform sulcus (MFS). LOTC was defined as previously described in Weiner and Grill-Spector (2013). Posteriorly, the LOTC ROI was defined at the convergence of the intraparietal sulcus (IPS) and the descending limb of the superior temporal sulcus (STS). The superior boundary was defined at the dorsal lip of the STS, and inferior boundary at the occipitotemporal sulcus (OTS). We then computed general linear models for all three individual fLoc localizer runs we acquired and computed *t*-statistic contrast maps identical to those used for our ROI definitions (e.g. faces vs all other categories, see ROI definition section for details), resulting in 3 contrast maps for each subject for each of the 4 categories: characters, bodies, faces, and places. Consequently, we computed the Dice coefficient between each pair of runs for each subject, hemisphere, and ROI, separately (run 1 and 2, 1 and 3, and 2 and 3 within VTC and LOTC). We then took the average across those three splits as the Dice coefficient for that subject. Ultimately, we performed this analysis with a liberal statistical threshold of *t* > 0 (any vertex holding a positive t-value is included) and once with a threshold of *t* = 2.2 (*p*<0.01) for vertices to be included in the contrast map (Fig. 4). Together, these measures result in a lower and upper bound estimation of our Dice coefficient noise ceiling.

**Figure 4.**
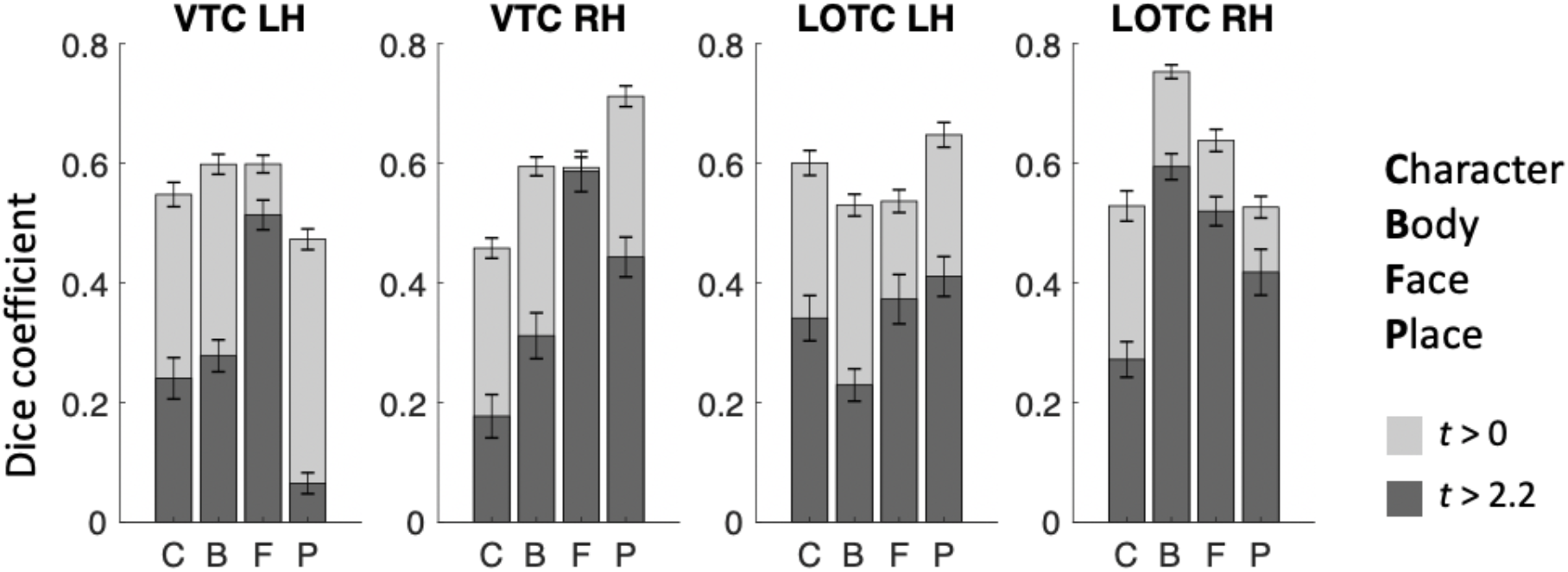
Reproducibility of category-selectivity responses. For the two cortical expanses that contain the category-selecitve regions of the visfAtlas, VTC and LOTC, the reproducibility of category responses was computed across the *t*-contrast maps of single runs for each respective category (see Materials and Methods for details). Dark gray bars represent the Dice coefficient results based on *t*-contrast maps that were thresholded with *t* > 2.2, which equals p < 0.01, while light gray bars were based on *t*-contrast maps that were thresholded at *t* > 0. Errorbars represent standard errors across subjects.

### Validation of the visfAtlas with an independent dataset of category-selectivity in ventral temporal cortex and with an increasing number of subjects

Common consideration in building atlases are (i) the number of subjects that are used to build the atlas and (ii) how well it can predict new datasets. To address whether our sample size is sufficient to achieve generalizability to new data, we tested how well the visfAtlas predicts fROIs of 12 new subjects. These data were acquired using a similar localizer in a different scanning facility, identified by independent experiments, and have been published previously (Stigliani et al. 2015; Weiner et al. 2017). We compared their fROI definitions of mFus-faces, pFus-faces, OTS-bodies, pOTS-characters and CoS-places to our visfAtlas definitions in the following ways: (1) We visualized our visfAtlas MPMs in relation to their probability maps of each of the fROIs (Fig. 6), and (2) we calculated how well our visfAtlas predicted each of their individual subjects’ fROIs using the Dice coefficient.

Lastly, to address how the number of subjects affects the accuracy of our visfAtlas, we calculated the Dice coefficient for different iterations of the visfAtlas in which we incrementally increased the number of subjects from 2 to 19; specifics are in the Supplemental Materials.

### Functional responses of atlas fROIs in left out data

When using a probabilistic atlas, it is of great interest not only to know how likely one would find a new subject’s fROI in the same location, but also what signals would be picked up for that subject within an atlas-fROI. For example, are voxel in face-selective atlas fROIs responding mostly to faces? To test the generalizability of our atlas, we performed a leave-subject-out maximum responsivity analysis. The analysis calculates the percentage of voxel responding highest to each condition within a given fROI, where the fROI is defined on all subject’s data except the one dataset used for the responsivity computation. This was repeated for all possible leave-subject-out combinations. First, for each subject individually we created a maximum probability map (MPM) based on the other N-1 subjects (leaving the target subject out). Then, for each individual voxel within each fROI in this MPM, we estimated the average response amplitude to each category across trials using the optimized Least Squares – Separate (LS-S) trial estimation approach as described by Mumford et al. (2012). Then, we created a ‘winner map’ for each fROI per subject, in which the condition index that yielded the strongest response was assigned to each voxel within the fROI. Per condition, we counted the number of winning voxels within the ROI, which we expressed as a percentage of the total number of voxels in the fROI. This procedure was repeated for each subject (Fig. 7).

### Comparison of our visfAtlas to existing publicly available atlases and relevant fROIs

How does the visfAtlas compare to published atlases? While there is no complete occipitotemporal atlas of visual areas yet, atlases of retinotopic areas have been published by Wang et al. (2014) and Benson et al. (2012, 2014). To compare our atlas to the Benson atlas where there is no separation between ventral and dorsal quarterfields, we merged our dorsal and ventral V1-V3. Additionally, there is a published probabilistic atlas of CoS-places (Weiner et al. 2018), and motion selective hMT+ (Huang et al. 2019). We compared our surface visfAtlas to the existing surface maps by assessing their correspondence in the FreeSurfer fsaverage space. For each published atlas we (i) qualitatively assessed the spatial correspondence by visualizing the atlas definitions on a common brain space of the FreeSurfer average brain (Fig. 8) and (ii) quantitatively assessed the correspondence by calculating the Dice coefficient between each of our individual subject’s fROIs and the respective other atlas as we do not have access to the individual subject data in the Wang, Benson or Huang atlases.

## RESULTS

Using data from 19 healthy participants we aimed at generating a probabilistic atlas of occipito-temporal and ventral temporal cortex. Individually defined regions were normalized to group space using either (1) cortex-based alignment (CBA) or (2) nonlinear volumetric alignment (NVA).

### Superior spatial overlap after cortex-based alignment for retinotopic and category selective regions

In order to determine whether nonlinear volumetric (NVA) or cortex-based alignment (CBA) result in higher accuracy and predictibility of our atlas, we aimed at comparing both alignment techniques across all functional regions of interest (fROIs). Figure 1 displays three example regions, one early visual retinotopic region in occipital cortex (V1d), as well as two higher-order category-selective regions in ventral temporal cortex (CoS-bodies and mFus-faces). Qualitatively, a higher degree of consistency across subjects is observable when group maps were normalized using CBA as compared to NVA. Both V1d and Cos-places display a high consistency in the group map center as indicated by yellow colored vertices, while centers are more variable after NVA alignment, most evident in V1d. For mFus-faces, both group maps display a greater degree of variability across subjects than the other two regions.

To quantify which group alignment resulted in higher consistency and therewith predictability, we used the Dice coefficent (DSC) and a leave-one-out cross-validation (LOOCV) procedure to determine the predictability of finding the same region in a new subject. Moreover, we calculated the Dice coefficient using different thresholds for the probabilistic group map, ranging from a liberal unthreshold (one subject at a given voxel/vertex is enough to assign it to the group map) map to a conservative threshold where all N-1 subjects had to share a voxel/vertex to be assigned to the group map (Fig. 2). For retinotopically defined regions, DSC’s varied between 0.35 and 0.59 for peak probability after CBA, and between 0.30 and 0.42 after NVA. Especially regions with a lower predictability overall tended to show higher predictability after NVA for more conservative group thresholds (e.g. Fig. 2B, mFus-faces, TOS-bodies). For CBA, peak predictibility (DSC) for each region ranged from 0.1 to 0.60, while it ranged from 0.1 to 0.42 for NVA, with character-selective regions showing the lowest consistency for both alignments, closely followed by mFus-and IOG-faces.

Quantitatively, CBA displayed an overall greater predictability across regions and thresholds (except for V3d LH, see Fig. 2A), which was confirmed by a significant difference in alignment for both unthresholded (F(1,34) = 20.12, *p* < .001) and thresholded (0.2; F(1,34) = 174.84, *p* < .001) probability maps, see Methods for details on threshold selection. Additionally, there was no significant main effect for hemisphere (unthresholded: *p* = .90; thresholded: *p* = .56) and no interaction between alignment and hemisphere (unthresholded: F(1,34) = .85, *p* = .36, thresholded: F(1,34) = 0.35, *p* = .56). We followed up with a paired permutation test (across alignments) for the unthresholded DSC within each fROI. As there was no main effect for hemisphere (see above) and no significant difference in region size across hemispheres (t(17) = - 0.48, *p* = .64, Fig. 3), permutation tests were performed on Dice coefficients using an unthresholded group map prediction and averaged across hemispheres. Results show that CBA alignment has a higher predictability than NVA for all regions (*p* < .05), except for unthresholded: pOTS-characters (*p* = 1), IOS-characters (*p* = .81), v3d (*p* = .05), IOG-faces (*p* = .05) and thresholded: V3d (*p* = .05), mFus (*p* = .70), IOG (*p* = .55), pOTS (*p* = 1), IOS (*p* = 1), OTS (*p* = .14).

As shown in the previous section and displayed in Figure 2, different category-selective regions in VTC and LOTC show different levels of Dice coefficients. One factor that may contribute to this variability is the region’s size, which also varies across fROIs (Fig. 3). To test if this relationship is significant, we measured the correlatation between the Dice coefficient and surface area of the fROIs. Results indicate a significant correaltion (left hemisphere: r = 0.83, *p* < 0.01; right hemisphere: r = 0.85, *p* < 0.01), suggesting that larger regions have higher Dice coefficients. We also examined if differences in Dice coefficient are related to differences in noise ceiling across ROIs. As a measure of noise ceiling, we calculated the within-subject Dice coefficient across the 3 runs of the fLoc. We reasoned that if there are between-ROIs differences in the noise ceiling estimated from within-subject Dice coefficients, they would also translate to the between-subject Dice coeffient. When using a lenient t-map threshold, results (Fig. 4) indicate that within-subject Dice coefficient for a lenient t-map threshold (t>0) range from 0.4 – 0.77 across categories. We find a higher Dice coefficient for bodies and faces in left VTC, and a higher Dice coefficient for places in the right VTC. In LOTC, the highest within-subject Dice coefficient is for place-seletivity in the left LOTC, and body-selecitivty in the right LOTC. Given that within- and between-subjects Dice coefficients are in the same range and vary similarly across fROIs, we believe that the precision of the visfAtlas will allow to identify fROIs in individual participants.

### A functional atlas of occipito-temporal cortex in volume and surface space

By systematically varying the group map threshold for predicting a left-out subject’s fROI, we established that a group map threshold of 0.2 allows for greatest predictability across regions. Using the 0.2 threshold, we generated a functional atlas of occipito-temporal cortex by generating a maximum probability map (MPM, see Methods for details). Figure 5 displays the resulting unique tiling of category-selective regions in stereotaxic space for surface (Fig. 5A) and volume (Fig. 5B) space. The visfAtlas is publicly available in both surface as well as volume space to allow usage in a variety of analyses and in file formats for BrainVoyager and FreeSurfer for surface space as well as in volume space using the NifTi format. In addition, we publish a BrainVoyager average brain (BVaverage, Fig. 5C; download.brainvoyager.com/data/visfAtlas.zip).

**Figure 5.**
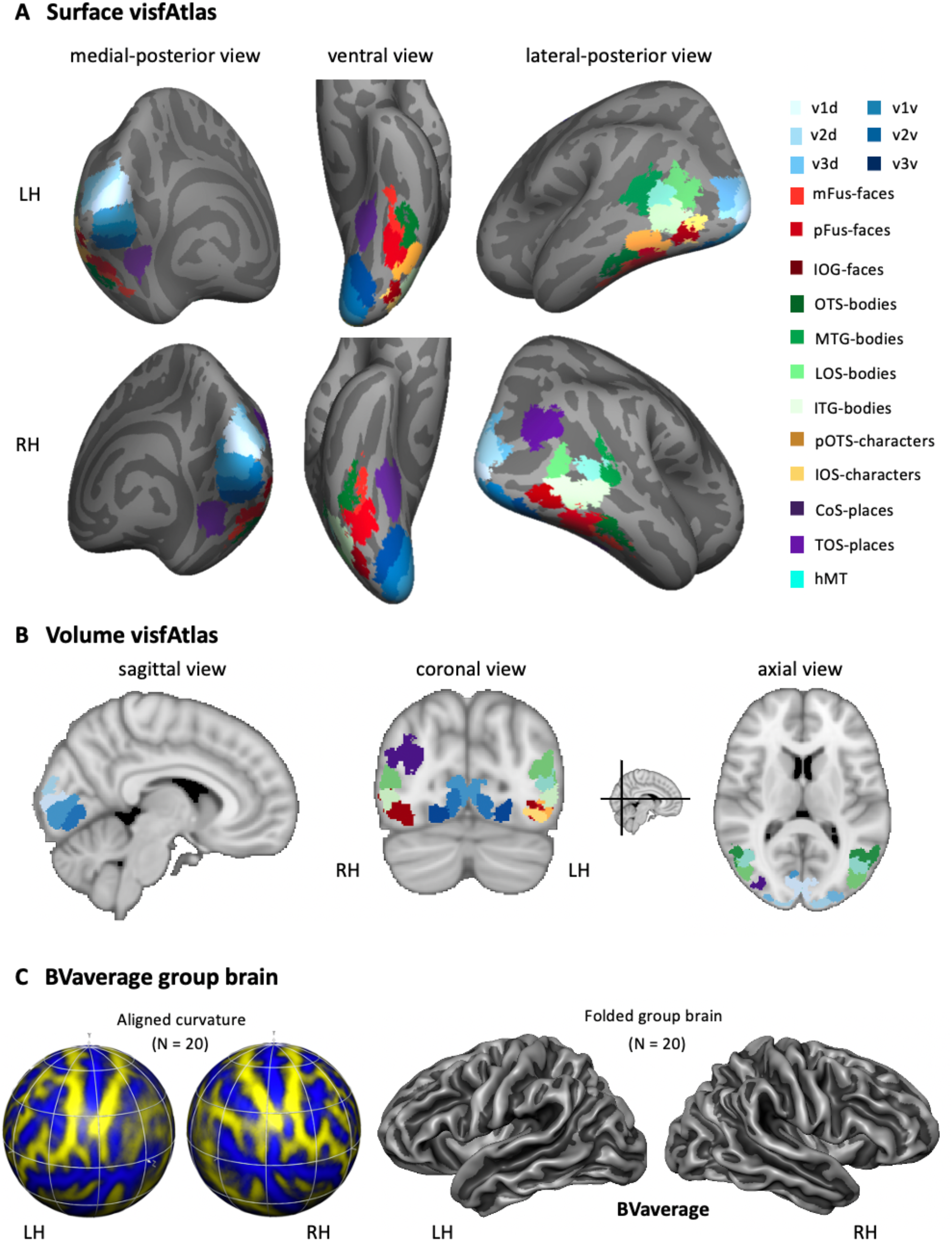
Maximum-probability map (MPM) of occipito-temporal cortex functional regions-of-interest (fROIs). (A) visfAtlas in surface space after cortex-based alignment. Each color displays a unique fROI group map thresholded at 0.2 of all subjects in which the given fROI could be identified. (B) Volume atlas using the same color coding as in surface space. Inset between coronal and axial view displays the slice location for coronal and axial slices, respectively. *LH*: left hemisphere, *RH*: right hemisphere. (C) A new group average brain (BVaverage) published in BrainVoyager, based on 20 adults. This average brain can be used for future studies as a common reference brain.

### Atlas validation using an independent dataset and an increasing number of subjects

How well does the visfAtlas localize regions in new subjects scanned at a different scanner and facility? To answer this question, we compared the ventral visfAtlas ROIs with a dataset acquired at Stanford University (Stigliani et al. 2015; Weiner et al. 2017) using different subjects and a functional localizer experiment similar to ours. Figure 6 shows unthresholded probabilistic maps of Weiner’s MPMs (across 12 participants) and our respective visfAtlas MPMs. Qualitatively, the location of their probabilistic maps, especially peak probabilities, correspond to our respective visfAtlas ROIs. To quantify the similarity, we tested how well our data predict the fROIs of these 12 independent subjects by calculating the Dice coefficient between our MPM fROIs and each of the independent subjects’ fROIs (Fig. 6B). The mean Dice coefficients (+/- SE) for left and right hemispheres, respectively, are in a similar range as the Dice coefficient of the leave-one-out-cross-validation results of our data (compare Fig. 2 threshold 0.2 with Fig. 6B).

**Figure 6.**
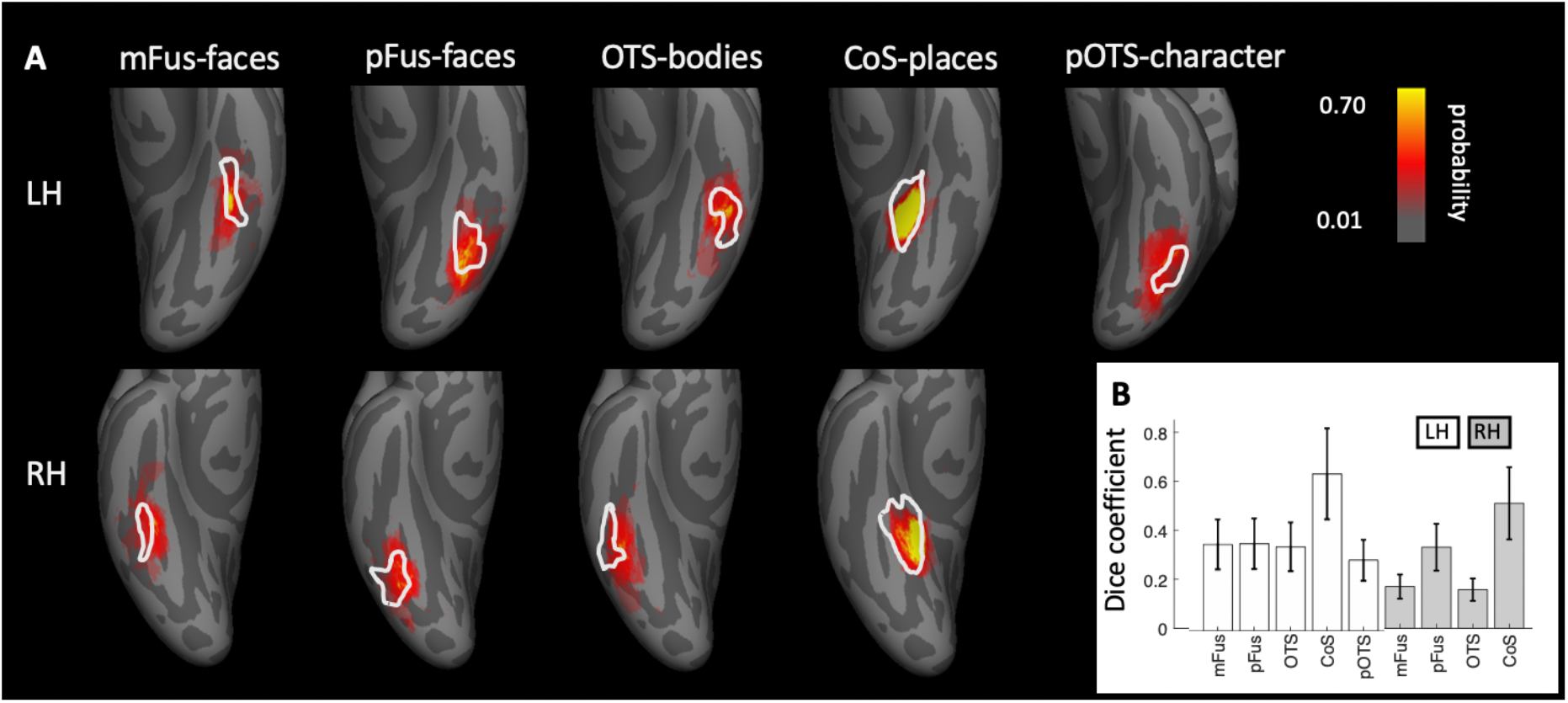
Correspondence between the visfAtlas and 5 ventral temporal cortex probabilistic maps from independent data. **(A)** We compared visfAtlas MPM fROIs (white outlines) in VTC with probabilistic maps (colored regions) of 6 functional regions from an independent dataset that used a similar localizer, which has been published previously (Stigliani et al. 2015; Weiner et al. 2017). *Top:* left hemisphere; *Bottom:* right hemisphere. **(B)** Average Dice coefficient between fROIs of the individual subjects from Stigliani and Weiner and colleagues and the MPMs of our visfAtlas fROIs. *Errorbars*: standard errors across subjects. *LH:* left hemisphere; *RH:* right hemisphere.

Additionally, we explored how the number of subjects used for generating our atlas affects its accuracy(Supplemental Fig. 1). Results indicate that in general, having more participants generates better accuracy in the LOOCV, but the number of requireed subjects varies across ROIs. Overall, across all ROIs, the highest Dice coefficient plateaus between 12 and 14 subjects, suggesting that our atlas based on an average of 16 subjects per ROI (see Table 1 for details) is sufficient.

### Generalizability of functional atlas: functional responsivity in left out data

One of the advantages of a probabilistic atlas is the ability to locate a region of interest with a degree of certainty (as established using the Dice coefficient analysis) in a new subject without the need to run a localizer itself. In order to quantify the atlas’ generalizability, the category responsivity of the category selective areas in new participants is a crucial metric. Therefore, we performed a leave-subject-out responsivity analysis in volume space to assess category-responsivity. For each fROI, we established the percentage of voxel that showed the strongest response to each available category (Fig. 7, see Methods for details of responsivity estimation). For all category selective regions, we confirmed that the category it is selective for indeed yields the highest percentage of maximum voxel responsivity across subjects. Face-selective fROIs (Fig. 7, top left) contain 52-72% (lowest to highest fROI) face-selective voxel responses (*red*). The second-highest maximum responsivity is body-selective (*green*) with 10-43% on average across subjects, followed by character-selective regions (*gray*) with 2-25%. Body-selective regions (Fig. 7, top right) contain the highest proportion of body as maximum voxel responsivity for lateral body-selective regions (80-94%), with lowest proportions for ventral OTS-bodies in left and right hemisphere (46-55%). The second-largest number of voxel-maximum-responsivity is faces (1-40%). Place-selective fROIs (Fig. 7, bottom left) show a large proportion of voxels with their preferred place responses (*purple*, 77-82%), followed by up to 21% body-maximum voxel responsivity. Character-selective ROIs (Fig 5., bottom right) on the other hand contain 41 - 52% character-response voxel, followed by up to 38% body-response voxels.

**Figure 7.**
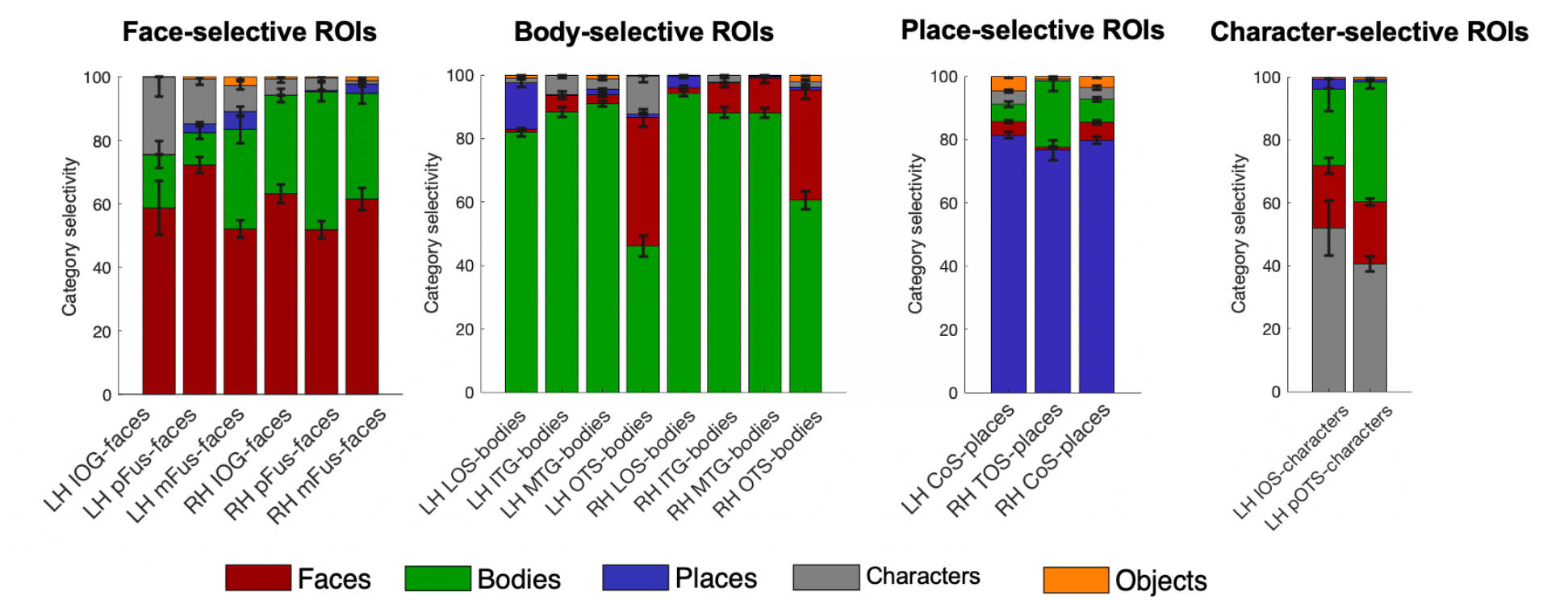
Proportion of voxels that show maximum responsivity in left out subjects are largely their own category. Using our volumetric atlas data we generated a cross-validated estimate of voxel maximum responsivity in a left out subject. N-1 times, we generated a volumetric maximum probability map and calculated the proportion of voxel that were maximally responsive for the ROI’s category, e.g. face response voxel in mFus-faces. This gives an estimate for the expected specificity of the atlas. For each major category - faces, bodies, places, characters – proportions of category responsivity are displayed with each region’s preferred category as the bottom bar of each stacked bar graph. *Error bars*: Proportion own category selectivity across all left-out subjects.

### Similarities between previously published atlas areas and our visfAtlas

In order to establish the correspondence of our probabilistic functional atlas to other atlases, we made quantitative comparisons to existing atlases of one or multiple regions localized with comparable stimuli. As retinotopic atlases are frequently used to define early visual cortices in new subjects, we wanted to compare our retinotopic areas V1-V3 dorsal and ventral to a group atlas of retinotopic visual areas aligned to the fsaverage brain by Wang et al. (2014). To assess the correspondence between the two atlases we computed the Dice coefficient (see Methods for details) between the existing group atlas and each of our visfAtlas subjects (Fig. 8) separately. Qualitatively, V1d and V1v from both atlases show a high degree of overlap and correspondence decreases when moving to the dorsal and ventral V2 and V3 (Fig. 8A). However, for each of the probabilistic maps of our visfAtlas regions, the peak probability location falls within the MPM published by Wang et al. (2014). This observation is confirmed by high Dice coefficients for V1d and V1v in the left and right hemisphere (average Dice coefficient 0.4 – 0.5, see Fig. 8E), and lower Dice coefficients in V2 and V3 (average Dice coefficient 0.15 – 0.4, Fig. 8E). Next, we also compared our visfAtlas retinotopic regions to an anatomical prediction of V1-V3 by Benson et al. (2012), which shows a similar pattern of correspondence with a greater overlap in V1 (0.4 – 0.42) and a decrease in V2 and V3 (0.2 – 0.29).

**Figure 8.**
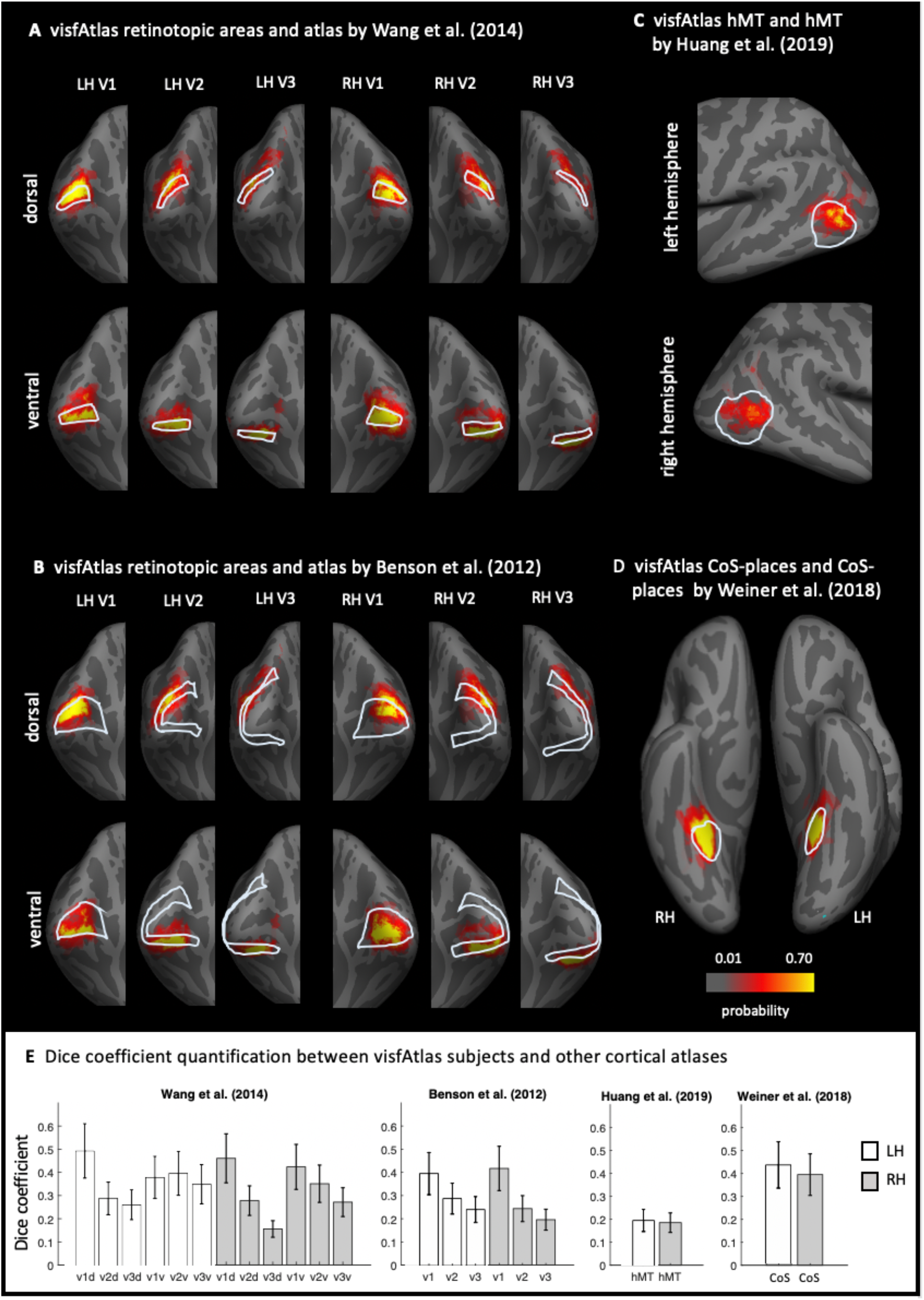
Comparison of the visfAtlas to other probabilistic atlases. In A-D each red-yellow map is the probabilistic map of unthresholded individual regions of the visfAtlas ROI and the outline is the fROI of the relevant atlas; all images are show in the fsaverage brain. (A) Comparison of V1-V3 dorsal and ventral of the retinotopic atlas published by Wang et al. (2014) and our respective visfAtlas regions. Regions are presented on a medial-occipital view of the fsaverage group brain. (B) Comparison of V1-V3 dorsal and ventral to the anatomically estimated V1-V3 (Benson et al. 2012). (C) Comparison of motion-selective hMT+ published by Huang et al. (2019) to visfAtlas hMT+ probabilistic map. (D) Comparison of CoS-places published by Weiner et al. (2018) to the visfAtlas CoS-places map. (E) Dice coefficient between the visfAtlas fROI and the same fROI defined by other atlases. *Errorbars:* Standard error across 19 visfAtlas subjects. LH: left hemisphere; *RH:* right hemisphere.

Similiar to the retinotopic regions, we compared a category-selective region - the CoS-places fROI - to a published probabilistic version by Weiner et al. (2018) which used a very similar localizer for their study. Both atlases display a high correspondence, with a slightly higher Dice coefficient in the left hemisphere than in the right hemisphere (Fig. 8E). On lateral occipito-temporal cortex we compared a recently published motion selective group area of hMT+ that has been defined using data from 509 adults (Huang et al. 2019).As Huang’s et al (2019) group fROI was not bounded by body-selective regions but ours was defined by maximum probability map (MPM) that takes into account the neighboring face and body-part areas, the visfAtlas is smaller than Huang’s definition Nonetheless, also here, the locus of our hMT+ probabilistic map is within the hMT+ atlas published by Huang et al (2019).

## DISCUSSION

In the present study, we generated a cross-validated functional atlas of occipito-temporal visual cortex, including early-visual cortex retinotopic regions as well as category-selective regions. Additionally, we evaluated how accurately this atlas predicts category-selectivity in left-out subjects. We found that cortex-based alignment (CBA) outperforms nonlinear volumetric alignment (NVA) for most ROIs. Importantly, using CBA our probabilistic category-selective ROIs accurately identify 40% - 94% of category-selective voxels in left-out subjects (Fig. 7). We make this functional atlas (visfatlas) of occipito-temporal cortex available on cortical surfaces of the fsaverage (FreeSurfer) and BVaverage (BrainVoyager), and volume formats in MNI space compatible with the majority of software tools.

In the following we will discuss the implications of our results for theories of anatomical and functional coupling in visual cortex, how our atlas relates to other atlases in the field, whether it can be validated by independent data, and how future research can expand on our atlas with new methodological approaches.

### Cortex-based alignment improves the consistency of group fROIs: Implications

Spatial consistency in both retinotopic and category-selective regions was on average higher after CBA as compared to NVA (Fig. 2). The higher performance of CBA is in agreement with previous studies that reported that CBA results in atlases with higher accuracy than volumetric atlases (Frost and Goebel 2012; Coalson et al. 2018), and specifically of retinotopic visual areas (Wang et al., 2014, Benson 2012) and cytoarchitectonic regions (Rosenke et al. 2017, 2018). Since CBA specifically aligns macroanatomical landmarks, the higher accuracy of CBA suggests a coupling between macroanatomical landmarks and functional regions. These results are consistent with prior research showing striking functional-macroanatomical coupling in visual cortex including: (i) V1 with the calcarine sulcus (Hinds et al. 2008), (ii) V3A and the transverse occipital sulcus (Nasr et al., 2011; Tootell et al., 1997), (iii) hV4 and the posterior transverse collateral sulcus (Witthoft et al., 2014), (iv) motion-selective hMT+ and the posterior inferior temporal sulcus (Dumoulin et al. 2000; Weiner and Grill-Spector 2011), (v) mFus-faces and the mid-fusiform sulcus (Grill-Spector and Weiner 2014) and (vi) CoS-places and the intersection of the anterior lingual sulcus with the collateral sulcus (Weiner et al. 2018). One interesting observation regarding the Dice coefficient results (Fig. 2) is that in some fROIs, NVA produces a higher Dice coefficient than CBA for high threshold values (e.g., pOTS-characters LH, mFus-faces RH). We hypothesize that since NVA is operating in 3D volume space and CBA in cortical surface space, shifts around crowns of gyri or fundi of sulci may produce a large impact on CBA than NVA. This hypothesis can be tested in future research.

Historically, the prevailing view (Glasser and Van Essen 2011; Haxby et al. 2011; Orban et al. 2014; Osher et al. 2015) was that higher-level functional visual regions have greater variability across participants as well as relative to macroanatomical landmarks compared to early visual areas such as V2 and V3. However, as we summarize in the prior paragraph, improvements in measurements and analysis methods argue against this prevailing view. In fact, our leave-one-out cross-validation procedure shows that five high-level visual regions (pFus-faces, LOS-bodies, ITG-bodies, CoS-places, motion-selective hMT+) have similar correspondence across subjects comparable to early visual cortex. However, some functional regions (mFus-faces, pOTS-characters, MTG-bodies, Fig. 2, see also Frost and Goebel, 2012), show more variability across participants. This diversity suggests that other factors may affect our ability to predict high-level visual regions. First, the shape and size of the ROI may impact across-subject alignment. Indeed, we found that larger and more convex ROIs tend to align better across participants than smaller ROIs, reflected in the finding of a positive correlation between the Dice coefficient and the size of the fROI. Second, the degree of macroanatomical variability differs across anatomical landmarks. In other words, stable macroanatomical landmarks may be better predictors of functional ROIs than variable ones. For example, the anterior tip of the mid-fusiform sulcus (MFS) is a more stable anatomical landmark than its posterior tip, as the length of the MFS substantially varies across people. Consequently, the anterior tip of the MFS better predicts face-selective mFus-faces than the posterior tip predicts pFus-faces (Weiner et al. 2014). Third, the quality of cortex-based alignment may vary across cortical locations (see Frost and Goebel 2012, 2013). Thus, more fragmented and less salient macroanatomical landmarks, such as the partially fragmented occipito-temporal sulcus (OTS), may align less well across participants with CBA. This in turn impacts the registration of functional ROIs that are associated with these landmarks. Fourth, the reliability of functional ROIs across sessions within an individual, which indicates a noise ceiling, may vary across ROIs. To evaluate the latter, we performed a reproducibility analysis for our category-selective regions by analyzing all three localizer runs independently (Fig. 4). This analysis highlights that running the same experiment multiple times within the same subject will not result in the exact same cortical activation pattern. Here, reproducibility estimates (Dice coefficients) ranged between 0.4 and 0.75 in VTC as well as LOTC, similar to Dice coefficient estimates by other studies (Weiner and Grill-Spector 2010; Weiner et al. 2016; Bugatus et al. 2017). Notably, the reproducibility analysis together with the analysis of an independent dataset indicate that reproducibility and variability of our Dice coefficient are within the range expected by previous studies (Weiner et al. 2018). However, one has to note that our reproducibility estimation is conservative since we used the three runs that comprised our category-selectivity localizer individually, which means that each split had less trials and a lower signal-to-noise ratio (SNR) than the analysis used to establish between-subject variability (3 runs per subject each). Future work should run the same experiment for an additional full 3 runs to establish a noise ceiling that is not impacted by SNR and trial number differences.

Future research can also improve the inter-subject alignment by improving CBA methods. For example, CBA may be improved by weighting microanatomical landmarks by their consistency and saliency. Other directions for improving the predictions of the model may include incorporating additional features, such as spatial relationships between ROIs, or adding some functional data (Frost and Goebel 2013) to improve predictions. For example, adding one retinotopic run improves predicting early visual areas relative to macroanatomical landmarks alone (Benson and Winawer 2018).

### Category-preferred responses within visfAtlas regions and reasons for variability across areas

As the main purpose of a functional atlas is to allow generalization to new individuals, confirmation and validation of the functional responses of the predicted regions is crucial. We used a leave-one-out-cross-validation approach to quantify the generalizability of our maximum probability map and demonstrate that voxels within the predicted ROI are displaying maximum responsivity to the preferred category of that ROI (Fig. 7). The highest proportion of own category-responsive voxels was in lateral body-selective regions and the lowest own category response was in character-selective regions. One possible explanation for this variability is the proximity of ROIs to regions selective for other categories. For example, in ventral temporal cortex, the body-selective region on the OTS is small and located between two larger face-selective regions, but in lateral occipito-temporal cortex, body-selective ROIs are larger and some of them distant from the face-selective regions on the IOG. Close proximity between ROIs selective for different categories increases the likelihood of overlapping atlas boundaries, which may reduce the predictions of category-selectivity in a new subject.

Another reason for variability across areas could be that areas are differentially affected by the number of subjects they require to reach a stable prediction. To test this, for each ROI we calculated Dice coefficients with N=2 to max N for that ROI and evaluated how the overlap changed with increasing number of subjects (Supplemental Fig. 1). Interestingly, our analysis suggests that not all ROIs benefit from an increasing number of subjects equally. More specifically, only 5 of the 18 ROIs displayed such an increase, and those suggest to plateau between 12 and 16 subjects. For other ROIs, the number of subjects did not impact the Dice coefficient. Generally, the assumption is that as the number of subjects increases, the level of noise decreases and one gets closer to the true between-subject variability. One interesting note is that using the data of our visfAtlas, none of the ROIs displayed a positive trend in Dice coefficient that continues past the number of subjects included in our atlas. Follow up work should evaluate whether this is local plateau or the global maximum Dice coefficient for each region.

Additionally, our approach can be extended to generate atlases of additional high-level visual regions that have other selectivities by including stimuli and contrasts for: (i) dynamic vs. still biological stimuli to identify regions selective for biological motion in the superior temporal sulcus (Puce et al. 1996; Grossman and Blake 2002; Beauchamp et al. 2003; Pitcher et al. 2011), (ii) objects vs. scrambled objects to identify object-selective regions of the lateral occipital complex (LOC; Malach et al. 1995; Grill-Spector et al. 1998; Vinberg and Grill-Spector 2008), and (iii) colored vs. black and white stimuli to identify color-selective regions in medial ventral temporal cortex (Beauchamp et al. 1999; Lafer-Sousa et al. 2016). Furthermore, future studies may explore the possibility to generate more sophisticated atlases, which contain not only a unique tiling of cortical regions, but also allow for multiple functional clusters to occupy overlapping areas and indicate probabilities for multiple categories at each voxel, perhaps building a hybrid of probabilistic maps of single regions and a maximum probability map.

### Consistent definitions of visual areas across different atlases

In generating our visfAtlas is was important for us to include early visual areas and hMT+ in addition to category-selective regions for two reasons: (1) it allowed us to benchmark and test our approach to atlases of retinotopic areas (e.g. Wang et al. 2014) and (2) it allowed us to generate a more comprehensive atlas of the visual system that includes the most studied visual regions spanning early and higher-level visual regions.

Finding that our approach generates similar ROIs to other atlases (e.g., V1-V3 in the Wang et al. (2014) atlas, Benson et al. (2012) atlas) and hMT+ (Huang et al. 2019) is important as it illustrates that these ROIs are robust to experimental design, stimuli type, and number of subjects that were used for generating atlases, all of which varied across studies. For example, we defined hMT+ by contrasting responses to expanding and contracting low contrast concentric rings to stationary ones in 19 subjects but Huang et al. (2019) defined hMT+ by contrasting responses to dots moving in several directions vs. stationary dots in 509 subjects. Despite these differences, where hMT+ is predicted to be, largely corresponds across both studies (Fig. 8C), even as the predicted spatial extend of hMT+ is substantially smaller in our atlas as compared to Huang’s. For retinotopic regions, we found the best correspondence between our data and Wang et al. (2014) for V1d and V1v, especially in the left hemisphere (Fig. 8A). Right hemisphere V1 of our visfAtlas extends more dorsally compared to Wang’s atlas, consequently shifting right hemisphere V2d and V3d further compared to Wang et al. (2014). For both, the comparison to Benson et al. (2012) and Wang et al. (2014), we observe a reduction in overlap that corresponds to a reduction in Dice coefficient when quantifying V1 vs. V2 and V3 (see Fig. 2 for details), indicating that these may be individual differences across subjects that are independent of anatomical coupling, but still display less individual variability than previously assumed (see Discussion section *Cortex-based alignment improves the consistency of group fROIs: Implications*).

Ultimately, the visfAtlas showed close correspondence to the comparison atlases, highlighting the robustness of our approach and the utility of functional atlases for future neuroimaging studies.

### Conclusion and future uses

To this date, no probabilistic atlas has been published which contains such an extensive set of functional regions in occipito-temporal cortex. The present study shows that most of the category-selective regions can be predicted in new subjects.

This functional atlas of occipito-temporal cortex is available in both surface and volume space and can be used in commonly used data formats such as BrainVoyager and FreeSurfer. We hope that this atlas may prove especially useful for (1) predicting a region of interest when no localizer data is available, saving scanning time and expenses, (2) comparisons across modalities and (3) patient populations, such as patients who have a brain lesion (Schiltz and Rossion 2006; Steeves et al. 2006; Sorger et al. 2007; Barton 2008; Gilaie-Dotan et al. 2009; de Heering and Rossion 2015) or are blind (Mahon et al. 2009; Bedny et al. 2011; Striem-Amit, Dakwar, et al. 2012; van den Hurk et al. 2017).

## Acknowledgements

We would like to thank Martin Frost for advice with the experimental design and Kevin S. Weiner for feedback on individual region definitions. RG was supported by the European FET Flagship project ‘Human Brain Project’ FP7-ICT-2013-FET-F/604102 Grant Agreements, No. 7202070 (SGA1) and No. 785907 (SGA2). KGS by NEI grant R01EY02391501.

## Supplemental Materials

**Supplemental Figure 1.**
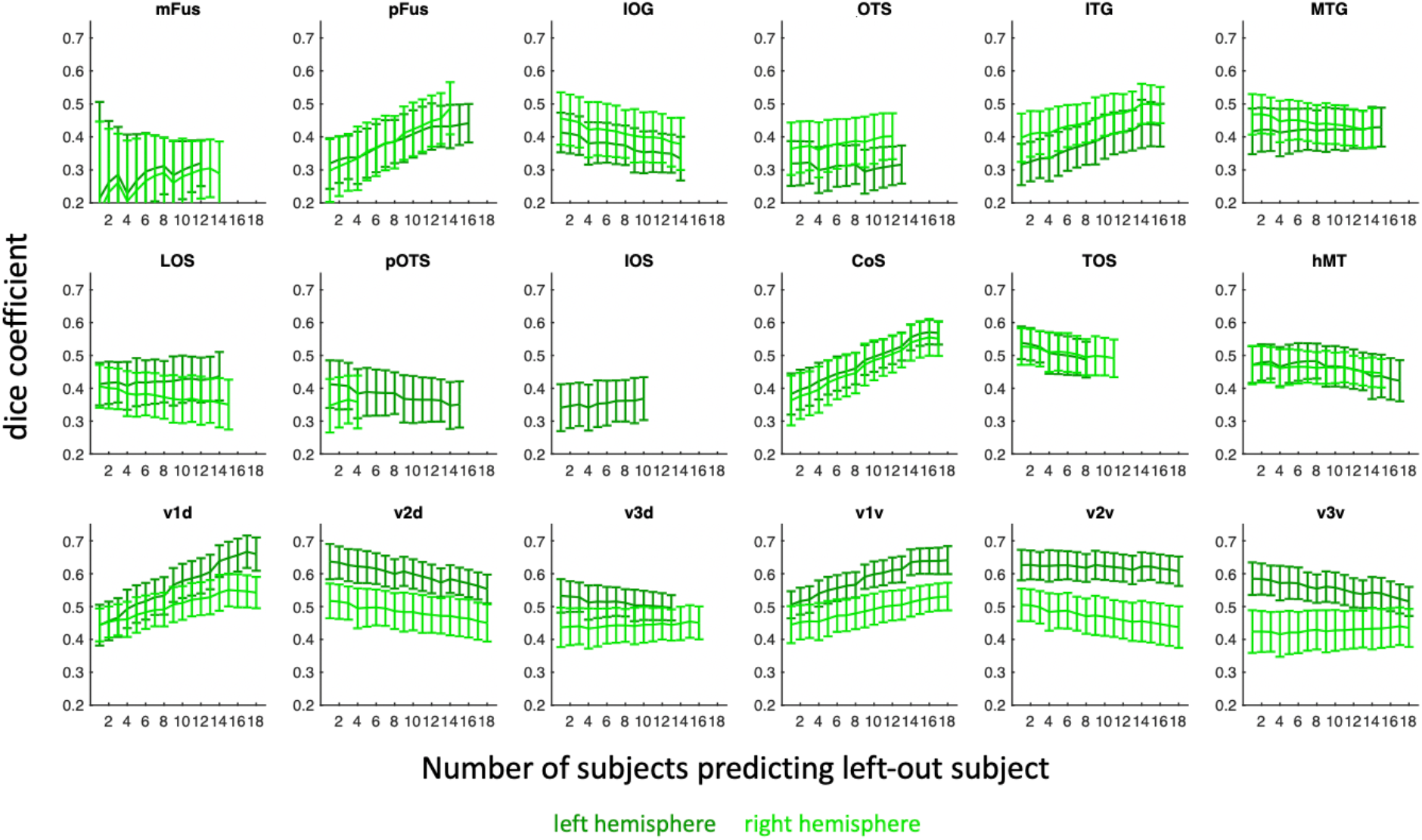
Effect of number of subjects in a group map on the Dice coefficient for predicting left-out subjects’ fROIs. For each iterative number of subjects comprising an atlas we tested how well it predicts a left-out data using the Dice coefficient metric. x-axis: number of subjects predicting a left-out subject; y-axis: resulting Dice coefficients. *Errorbars:* standard deviation across 1000 sample computations.

### Methodological approach

To evaluate the effect of number of subjects on the Dice coefficient for a given fROI, we calculated the Dice coefficient with an iterative number of subjects comprising the predicting group maps for each fROI in the visfAtlas. Details for the number of subjects that each fROI was defined in can be found in **Table 1** of the main article. For each fROI and hemisphere, respectively, we started with N = 2 subjects where 1 subject was used to predict the other subject. Then we randomly, without replacement, drew N = 3 subjects and used two to predict the third subject. The prediction was quantitively evaluated with the Dice coefficient. For any predicting number of subjects and each fROI, we used the same threshold that was best across all alignment methods (see main article, Methods and Materials), which was 0.2. Within each iteration of a given number of subjects, we cross-validated the dice coefficient for each left-out subject so that there were three different cross-validation iterations for N = 3, since each subject was left out once. The number of subjects was increased until the total N for the respective hemisphere ROI was reached (see x-axis of Suppl. Fig. 1). Next, we repeated this procedure for each N 1000 times (where the same subjects could not be drawn within the same sample, but could for any of the 1000 times) and computed the standard deviation across those iterations. We chose to draw samples 1000 times to control for the fact that there are more possible combinations of lower N than higher N.

## References

Aguirre GK, Zarahn E, D’Esposito M. 1998. An Area within Human Ventral Cortex Sensitive to “Building” Stimuli. Neuron. 21:373–383.

Amunts K, Malikovic A, Mohlberg H, Schormann T, Zilles K. 2000. Brodmann’s areas 17 and 18 brought into stereotaxic pace - where and how variable? . Neuroimage. 11:66–84.

Arcaro MJ, McMains SA, Singer BD, Kastner S. 2009. Retinotopic organization of human ventral visual cortex. J Neurosci. 29:10638–10652.

Barton JJS. 2008. Structure and function in acquired prosopagnosia: lessons from a series of 10 patients with brain damage. J Neuropsychol. 2:197.

Beauchamp MS, Haxby J V, Jennings JE, Deyoe EA. 1999. An fMRI Version of the Farnsworth – Munsell 100-Hue Test Reveals Multiple Color-selective Areas in Human Ventral Occipitotemporal Cortex. Cereb Cortex. 9:257–263.

Beauchamp MS, Lee KE, Haxby J V., Martin A. 2003. fMRI Responses to Video and Point-Light Displays of Moving Humans and Manipulable Objects. J Cogn Neurosci. 15:991–1001.

Bedny M, Pascual-Leone A, Dodell-Feder D, Fedorenko E, Saxe R. 2011. Language processing in the occipital cortex of congenitally blind adults. Proc Natl Acad Sci U S A. 108:4429– 4434.

Benson NC, Butt OH, Brainard DH, Aguirre GK. 2014. Correction of Distortion in Flattened Representations of the Cortical Surface Allows Prediction of V1-V3 Functional Organization from Anatomy. PLoS Comput Biol. 10.

Benson NC, Butt OH, Datta R, Radoeva PD, Brainard DH, Aguirre GK. 2012. The retinotopic organization of striate cortex is well predicted by surface topology. Curr Biol. 22:2081– 2085.

Benson NC, Winawer J. 2018. Bayesian analysis of retinotopic maps. Elife. 7:1–29.

Bugatus L, Weiner KS, Grill-Spector K. 2017. Task alters category representations in prefrontal but not high-level visual cortex. Neuroimage. 155:437–449.

Caspers J, Zilles K, Eickhoff SB, Schleicher A, Mohlberg H, Amunts K. 2013. Cytoarchitectonical analysis and probabilistic mapping of two extrastriate areas of the human posterior fusiform gyrus. Brain Struct Funct. 218:511–526.

Coalson TS, Van Essen DC, Glasser MF. 2018. The impact of traditional neuroimaging methods on the spatial localization of cortical areas. Proc Natl Acad Sci U S A. 115:E6356–E6365.

Cohen L, Dehaene S, Naccache L, Lehericy S, Dehaene-Lambertz G, Henaff M, Michel F. 2000. The visual word form area: Spatial and temporal characterization of an initial stage of reading in normal subjects and posterior split-brain patients. Brain. 123:291–307.

de Heering A, Rossion B. 2015. Rapid categorization of natural face images in the infant right hemisphere. Elife. 4:1–14.

DeYoe E a, Carman GJ, Bandettini P, Glickman S, Wieser J, Cox R, Miller D, Neitz J. 1996. Mapping striate and extrastriate visual areas in human cerebral cortex. Proc Natl Acad Sci U S A. 93:2382–2386.

Downing PE, Downing PE, Jiang Y, Jiang Y, Shuman M, Shuman M, Kanwisher N, Kanwisher N. 2001. A cortical area selective for visual processing of the human body. Science. 293:2470–2473.

Dumoulin SO, Bittar RG, Kabani NJ, Baker CL, Le Goualher G, Bruce Pike G, Evans a C. 2000. A new anatomical landmark for reliable identification of human area V5/MT: a quantitative analysis of sulcal patterning. Cereb Cortex. 10:454–463.

Dumoulin SO, Wandell BA. 2008. Population receptive field estimates in human visual cortex. Neuroimage. 39:647–660.

Eickhoff SB, Stephan KE, Mohlberg H, Grefkes C, Fink GR, Amunts K, Zilles K. 2005. A new SPM toolbox for combining probabilistic cytoarchitectonic maps and functional imaging data. Neuroimage. 25:1325–1335.

Emmerling TC, Zimmermann J, Sorger B, Frost MA, Goebel R. 2016. Decoding the direction of imagined visual motion using 7 T ultra-high field fMRI. Neuroimage. 125:61–73.

Engel SA, Glover GH, Wandell BA. 1997. Retinotopic organization in human visual cortex and the spatial precision of functional MRI. Cereb Cortex. 7:181–192.

Engel SA, Rumelhart DE, Wandell BA, Lee AT, Glover GH, Chichilnisky E-J, Shadlen MN. 1994. fMRI of human visual cortex. Nature.

Engell AD, McCarthy G. 2013. Probabilistic atlases for face and biological motion perception: An analysis of their reliability and overlap. Neuroimage. 74:140–151.

Epstein R, Kanwisher N. 1998. A cortical representation of the local visual environment. Nature. 392:598–601.

Frost MA, Goebel R. 2012. Measuring structural-functional correspondence: spatial variability of specialised brain regions after macro-anatomical alignment. Neuroimage. 59:1369–1381.

Frost MA, Goebel R. 2013. Functionally informed cortex based alignment: an integrated approach for whole-cortex macro-anatomical and ROI-based functional alignment. Neuroimage. 83:1002–1010.

Gilaie-Dotan S, Perry A, Bonneh Y, Malach R, Bentin S. 2009. Seeing with profoundly deactivated mid-level visual areas: Non-hierarchical functioning in the human visual cortex. Cereb Cortex. 19:1687–1703.

Glasser MF, Coalson TS, Robinson EC, Hacker CD, Harwell J, Yacoub E. 2016. A multi-modal parcellation of human cerebral cortex. Nat Publ Gr. 536:171–178.

Glasser MF, Van Essen DC. 2011. Mapping Human Cortical Areas In Vivo Based on Myelin Content as Revealed by T1- and T2-Weighted MRI. J Neurosci. 31:11597 LP – 11616.

Goebel R, Esposito F, Formisano E. 2006. Analysis of functional image analysis contest (FIAC) data with brainvoyager QX: From single-subject to cortically aligned group general linear model analysis and self-organizing group independent component analysis. Hum Brain Mapp. 27:392–401.

Grill-Spector K, Kushnir T, Edelman S, Itzchak Y, Malach R. 1998. Cue-invariant activation in object-related areas of the human occipital lobe. Neuron. 21:191–202.

Grill-Spector K, Weiner KS. 2014. The functional architecture of the ventral temporal cortex and its role in categorization. Nat Rev Neurosci. 15:536–548.

Grossman ED, Blake R. 2002. Brain areas active during visual perception of biological motion. Neuron. 35:1167–1175.

Hasson U, Harel M, Levy I, Malach R. 2003. Large-scale mirror-symmetry organization of human occipito-temporal object areas. Neuron. 37:1027–1041.

Haxby JV, Guntupalli JS, Connolly AC, Halchenko YO, Conroy BR, Gobbini MI, Hanke M, Ramadge PJ. 2011. A common, high-dimensional model of the representational space in human ventral temporal cortex. Neuron. 72:404–416.

Hinds OP, Rajendran N, Polimeni JR, Augustinack JC, Wiggins G, Wald LL, Diana Rosas H, Potthast A, Schwartz EL, Fischl B. 2008. Accurate prediction of V1 location from cortical folds in a surface coordinate system. Neuroimage. 39:1585–1599.

Huang T, Chen X, Jiang J, Zhen Z, Liu J. 2019. A probabilistic atlas of the human motion complex built from large-scale functional localizer data. Hum Brain Mapp. 40:hbm.24610.

Huk AC, Dougherty RF, Heeger DJ. 2002. Retinotopy and functional subdivision of human areas MT and MST. J Neurosci. 22:7195–7205.

Julian JB, Fedorenko E, Webster J, Kanwisher N. 2012. An algorithmic method for functionally defining regions of interest in the ventral visual pathway. Neuroimage. 60:2357–2364.

Kanwisher NG, McDermott J, Chun MM. 1997. The Fusiform Face Area: A Module in Human Extrastriate Cortex Specialized for Face Perception. J Neurosci. 17:4302–4311.

Kujovic M, Zilles K, Malikovic A, Schleicher A, Mohlberg H, Rottschy C, Eickhoff SB, Amunts K. 2013. Cytoarchitectonic mapping of the human dorsal extrastriate cortex. Brain Struct Funct. 218:157–172.

Lafer-Sousa R, Conway BR, Kanwisher NG. 2016. Color-Biased Regions of the Ventral Visual Pathway Lie between Face- and Place-Selective Regions in Humans, as in Macaques. J Neurosci. 36:1682–1697.

Lorenz S, Weiner KS, Caspers J, Mohlberg H, Schleicher A, Bludau S, Eickhoff SB, Grill-Spector K, Zilles K, Amunts K. 2015. Two New Cytoarchitectonic Areas on the Human Mid-Fusiform Gyrus. Cereb Cortex. 1–13.

Mahon BZ, Anzellotti S, Schwarzbach J, Zampini M, Caramazza A. 2009. Category-Specific Organization in the Human Brain Does Not Require Visual Experience. Neuron. 63:397–405.

Malach R, Reppas JB, Benson RR, Kwong KK, Jiang H, Kennedy W a, Ledden PJ, Brady TJ, Rosen BR, Tootell RB. 1995. Object-related activity revealed by functional magnetic resonance imaging in human occipital cortex. Proc Natl Acad Sci U S A. 92:8135–8139.

Mumford JA, Turner BO, Ashby FG, Poldrack RA. 2012. Deconvolving BOLD activation in event-related designs for multivoxel pattern classification analyses. Neuroimage. 59:2636– 2643.

Nasr S, Liu N, Devaney KJ, Yue X, Rajimehr R, Ungerleider LG, Tootell RBH. 2011. Scene-selective cortical regions in human and nonhuman primates. J Neurosci. 31:13771–13785.

Nieto-Castañón A, Fedorenko E. 2012. Subject-specific functional localizers increase sensitivity and functional resolution of multi-subject analyses. Neuroimage. 63:1646–1669.

Orban GA, Zhu Q, Vanduffel W. 2014. The transition in the ventral stream from feature to real-world entity representations. Front Psychol. 5:1–9.

Osher DE, Saxe RR, Koldewyn K, Gabrieli JDE, Kanwisher N, Saygin ZM. 2015. Structural Connectivity Fingerprints Predict Cortical Selectivity for Multiple Visual Categories across Cortex. Cereb Cortex. 1–16.

Peelen MV., Glaser B, Vuilleumier P, Eliez S. 2009. Differential development of selectivity for faces and bodies in the fusiform gyrus. Dev Sci. 12:16–25.

Peelen MV, Downing PE. 2005. Selectivity for the human body in the fusiform gyrus. J Neurophysiol. 93:603–608.

Pitcher D, Dilks DD, Saxe RR, Triantafyllou C, Kanwisher N. 2011. Differential selectivity for dynamic versus static information in face-selective cortical regions. Neuroimage. 56:2356–2363.

Puce A, Allison T, Asgari M, Gore JC, McCarthy G. 1996. Differential sensitivity of human visual cortex to faces, letterstrings, and textures: A functional magnetic resonance imaging study. J Neurosci. 16:5205–5215.

Rosenke M, Weiner KS, Barnett MA, Zilles K, Amunts K, Goebel R, Grill-Spector K. 2017. Data on a cytoarchitectonic brain atlas: effects of brain template and a comparison to a multimodal atlas. Data Br. 12:327–332.

Rosenke M, Weiner KS, Barnett MA, Zilles K, Amunts K, Goebel R, Grill-Spector K. 2018. A cross-validated cytoarchitectonic atlas of the human ventral visual stream. Neuroimage. 170:257–270.

Rottschy C, Eickhoff SB, Schleicher A, Mohlberg H, Kujovic M, Zilles K, Amunts K. 2007. Ventral visual cortex in humans: Cytoarchitectonic mapping of two extrastriate areas. Hum Brain Mapp. 28:1045–1059.

Saxe R, Brett M, Kanwisher N. 2006. Divide and conquer: a defense of functional localizers. Neuroimage. 30:1088–1089.

Schiltz C, Rossion B. 2006. Faces are represented holistically in the human occipito-temporal cortex. Neuroimage. 32:1385–1394.

Schwarzlose RF, Baker CI, Kanwisher N. 2005. Separate Face and Body Selectivity on the Fusiform Gyrus. J Neurosci. 25:11055–11059.

Senden M, Reithler J, Gijsen S, Goebel R. 2014. Evaluating Population Receptive Field Estimation Frameworks in Terms of Robustness and Reproducibility. PLoS One. 9:e114054.

Sereno MI, Dale a M, Reppas JB, Kwong KK, Belliveau JW, Brady TJ, Rosen BR, Tootell RBH, Series N, May N. 1995. Borders of Multiple Visual Areas in Humans Revealed by Functional Magnetic Resonance Imaging Borders of Multiple Visual Areas in Humans Revealed by Functional Magnetic Resonance Imaging. 268:889–893.

Sorger B, Goebel R, Schiltz C, Rossion B. 2007. Understanding the functional neuroanatomy of acquired prosopagnosia. Neuroimage. 35:836–852.

Steeves JKE, Culham JC, Duchaine BC, Pratesi CC, Valyear KF, Schindler I, Humphrey GK, Milner AD, Goodale MA. 2006. The fusiform face area is not sufficient for face recognition: Evidence from a patient with dense prosopagnosia and no occipital face area. Neuropsychologia. 44:594–609.

Stigliani A, Weiner KS, Grill-Spector K. 2015. Temporal Processing Capacity in High-Level Visual Cortex Is Domain Specific. J Neurosci. 35:12412–12424.

Striem-Amit E, Cohen L, Dehaene S, Amedi A. 2012. Reading with Sounds: Sensory Substitution Selectively Activates the Visual Word Form Area in the Blind. Neuron. 76:640–652.

Striem-Amit E, Dakwar O, Reich L, Amedi A. 2012. The large-scale organization of “visual” streams emerges without visual experience. Cereb Cortex. 22:1698–1709.

Susilo T, Yang H, Potter Z, Robbins R, Duchaine B. 2015. Normal Body Perception despite the Loss of Right Fusiform Gyrus. J Cogn Neurosci. 27:614–622.

Talairach J, Tournoux P. 1988. Co-planar Stereotaxic Atlas of the Human Brain.

Tootell RB, Mendola JD, Hadjikhani NK, Ledden PJ, Liu AK, Reppas JB, Sereno MI, Dale AM. 1997. Functional analysis of V3A and related areas in human visual cortex. J Neurosci. 17:7060–7078.

van den Hurk J, Van Baelen M, Op de Beeck HP. 2017. Development of visual category selectivity in ventral visual cortex does not require visual experience. Proc Natl Acad Sci. 114:E4501–E4510.

Vinberg J, Grill-Spector K. 2008. Representation of shapes, edges, and surfaces across multiple cues in the human visual cortex. J Neurophysiol.

Wandell BA, Brewer AA, Dougherty RF. 2005. Visual field map clusters in human cortex. Philos Trans R Soc Lond B Biol Sci. 360:693–707.

Wandell BA, Winawer J. 2011. Imaging retinotopic maps in the human brain. Vision Res. 51:718–737.

Wang L, Mruczek REB, Arcaro MJ, Kastner S. 2014. Probabilistic Maps of Visual Topography in Human Cortex. Cereb Cortex. 1–21.

Weiner KS, Barnett M, Lorenz S, Caspers J, Stigliani A, Amunts K, Zilles K, Fischl B, Grill-Spector K. 2017. The Cytoarchitecture of Domain-specific Regions in Human High-level Visual Cortex. Cereb Cortex. 1–16.

Weiner KS, Barnett MA, Witthoft N, Golarai G, Stigliani A, Kay KN, Gomez J, Natu VS, Amunts K, Zilles K, Grill-Spector K. 2018. Defining the most probable location of the parahippocampal place area using cortex-based alignment and cross-validation. Neuroimage. 170:373–384.

Weiner KS, Golarai G, Caspers J, Chuapoco MR, Mohlberg H, Zilles K, Amunts K, Grill-Spector K. 2014. The mid-fusiform sulcus: a landmark identifying both cytoarchitectonic and functional divisions of human ventral temporal cortex. Neuroimage. 84:453–465.

Weiner KS, Grill-Spector K. 2010. Sparsely-distributed organization of face and limb activations in human ventral temporal cortex. Neuroimage. 52:1559–1573.

Weiner KS, Grill-Spector K. 2011. Not one extrastriate body area: Using anatomical landmarks, hMT+, and visual field maps to parcellate limb-selective activations in human lateral occipitotemporal cortex. Neuroimage. 56:2183–2199.

Weiner KS, Grill-Spector K. 2013. Neural representations of faces and limbs neighbor in human high-level visual cortex: Evidence for a new organization principle. Psychol Res. 77:74–97.

Weiner KS, Jonas J, Gomez J, Maillard L, Brissart H, Hossu G, Jacques C, Loftus D, Colnat-Coulbois S, Stigliani A, Barnett MA, Grill-Spector K, Rossion B. 2016. The face-processing network is resilient to focal resection of human visual cortex. J Neurosci. 36:8425–8440.

Witthoft N, Nguyen M, Golarai G, LaRocque KF, Liberman A, Smith ME, Grill-Spector K. 2014. Where is human V4? Predicting the location of hV4 and VO1 from cortical folding. Cereb Cortex. 24:2401–2408.

Zeki S, Kennard C, Watson JDG, Lueck CJ, Frackowiak RSJ. 1991. Cortex of Functional Specialization in Human Visual. 17.

Zhen Z, Kong X-Z, Huang L, Yang Z, Wang X, Hao X, Huang T, Song Y, Liu J. 2017. Quantifying the variability of scene-selective regions: Interindividual, interhemispheric, and sex differences. Hum Brain Mapp. 38:2260–2275.

Zimmermann J, Goebel R, de Martino F, van de Moortele PF, Feinberg D, Adriany G, Chaimow D, Shmuel A, Uǧurbil K, Yacoub E. 2011. Mapping the organization of axis of motion selective features in human area mt using high-field fmri. PLoS One. 6:1–10.

